# Macrocyclic Peptides Containing an Imidazopyridinium (IP^+^) Unit Display Enhanced Passive Cell Permeability

**DOI:** 10.1101/2025.10.10.681645

**Authors:** Bo Li, Joshua Parker, Skyler Briggs, Jiajun Dong, James M. Burke, Thomas Kodadek

## Abstract

Macrocyclic peptides (MPs) have emerged as interesting therapeutic candidates due to their ability to engage difficult protein targets with high affinity and selectivity. However, their application to intracellular targets is limited by the poor passive membrane permeability of most MPs. We previously showed that incorporation of an imidazopyridinium (IP^+^) moiety into an MP significantly boosted passive membrane permeability, as measured by the parallel artificial membrane permeability assay (PAMPA)(Li, et al. (2024) *J. Amer. Chem. Soc*. **146**, 14633-14644). In this study we report a detailed analysis of the entry of IP^+^-containing MPs into living cells. Chloroalkane penetration assay (CAPA) data show that IP^+^ MPs access the cytoplasm rapidly, often at rates approaching those of drug-like small molecules. Mechanistic studies, including live-cell imaging, ATP-depletion experiments, and organelle colocalization analyses, indicate that IP^+^ MPs traverse the plasma membrane primarily via passive diffusion, avoiding endosomal entrapment. IP^+^ MPs do not localize to mitochondria, as is the case for many positively charged molecules. We show that incorporation of an IP^+^ unit transforms a previously described membrane impermeable macrocyclic antagonist of the p53–MDM2 interaction into a bioactive inhibitor of MCF-7 proliferation. Collectively, these results establish that IP^+^ incorporation is an effective strategy for the development of bioactive MPs targeting intracellular proteins.

## Introduction

Macrocyclic peptides (MPs) are currently of great interest to chemical biologists and medicinal chemists due to their ability to engage difficult protein targets with excellent affinity and selectivity. The discovery of orally bioavailable MPs MK-0616 ^1^ (a PCSK9 inhibitor), LUNA18^2^ (a Kras inhibitor), and Icotrokinra^3^ (an IL-23 Receptor antagonist), represent outstanding recent examples of the therapeutic potential of MPs. However, a persistent challenge remains: the inherently poor membrane permeability of most MPs. This is largely due to the fact that the N-H unit in the peptide bond is an excellent hydrogen bond donor, which results in it being well-hydrated in aqueous solution. This impedes passive diffusion of the MP through the plasma membrane. Indeed, one of Lipinski’s “rules” is that a drug candidate should have no more than five hydrogen bond donors.^4^

The vast majority of efforts to surmount this limitation have focused on emulating the various features of bioactive natural products such as Cyclosporine. These molecules display a high percentage of N-methylated amide bonds and are able to adopt conformations in an apolar environment that promote the formation of intramolecular hydrogen bonds that “hide” the remaining amide N-H units from water. In addition, cell permeable MP natural products almost exclusively contain hydrophobic side chains. Indeed, the ChemDraw-calculated logP (clogP) of Cyclosporine A is 14.4, though the actual logP is much lower than this value. The developers of LUNA18 at Chugai Pharmaceuticals, who have focused considerable effort on optimizing the passive permeability of MPs, suggest that a fruitful area of “chemical space” in which to search for cell-active MPs is described by eleven residues or less, with a clogP of 12.9 or higher and six or more N-methylated peptide bonds.^2^

While effective, strategies that focus primarily on increasing lipophilicity narrow the chemical space and structural diversity available for MPs drug discovery. Thus, new design principles are needed to broaden the scope of permeable MPs. Recently, we described the synthesis of novel MPs in which the ring is closed through the formation an imidazopyridinium (IP^+^)^5^ unit.^6^ We hypothesized that the permanent positive charge (i.e., not due to a protonation event) of this moiety would attract the IP^+^ MPs to the surface of a phospholipid bilayer after which the hydrophobic nature of the ring system might facilitate subsequent movement through the membrane. Encouragingly, we showed that many IP^+^ MPs with molecular weights (MWs) ranging from 600-1200 traversed the membrane with rates typical of much smaller, drug-like molecules in the parallel artificial membrane permeability assay (PAMPA).

While useful, the PAMPA employs an artificial membrane that does not reflect all of the properties of the plasma membrane of living cells.^7^ Therefore, in this study, we evaluate in detail the entry of IP^+^ MPs into mammalian cells. We find that in almost all cases, inclusion of an IP^+^ unit improves cytosolic exposure to an MP, in some cases by up to 40-fold. IP^+^ MPs are shown to immediately concentrate on the surface of the plasma membrane when introduced to mammalian cells, then slip rapidly into the cell within minutes. Several lines of evidence show that cell entry occurs by passive permeation through the membrane, not by endocytosis or some other active transport mechanism. Once inside the cell IP^+^ MPs, unlike some other positively charged molecules, do not localize to the mitochondrial membrane.

These data suggested that inclusion of an IP^+^ unit into existing MPs with good *in vitro* activity but poor permeability might stimulate their cellular activity. As a first test of this idea, we show that IP^+^ incorporation into a previously reported, cell impermeable MP antagonist of the p53-Mdm2 interaction^8^ dramatically improves the toxicity of the molecule to MCF-7 breast cancer cells. Taken together, the data reported here argue that IP^+^ incorporation into MPs provides a powerful tool for improving the passive cell permeability of this class of molecules that will expand the chemical space available for development of “beyond rule of 5” (Bro5) probe molecules and drug candidates targeting intracellular proteins.

## Results

### Synthesis of IP^+^ MPs with a Chloroalkane Tag (Ct)

The chloroalkane penetration assay (CAPA)^9^ provides a convenient method to measure the relative cell permeabilities of a series of molecules. The molecule of interest (MOI) is linked to a chloroalkane tag (Ct). Cells that express the HaloTag protein^10^ are exposed to a given concentration of this chimera for a certain period of time. After washing away excess MOI-Ct, the cells are then challenged with a fluorescent Ct conjugate. Any Halotag protein that has not been alkylated by the MOI-Ct will retain the fluorescent label. After another wash, the level of cellular fluorescence is measured. The permeability of the MOI-Ct conjugate is inversely proportional to the amount of fluorescence retained by the cells.

We first established an efficient solid-phase synthesis of IP^+^-linked macrocyclic peptides bearing the Ct (**Figure 1**). The synthetic route was fully compatible with Fmoc solid-phase peptide synthesis (SPPS) on TentaGel beads. Briefly, linear peptide aldehydes were assembled on resin by standard Fmoc chemistry followed by oxidation of the N-terminal serine residue.^11^ Macrocyclization was then achieved through condensation of the N-terminal aldehyde, a lysine side chain, and pyridine-2-carboxaldehyde (P2CA), thereby generating the IP^+^ unit.^6^ The Ct moiety was introduced via amide coupling with either amines or carboxylic acids positioned on the macrocycle. Finally, the peptides were cleaved from resin using trifluoroacetic acid (TFA), and purified by HPLC. Full experimental details are provided in the Supporting Information.

**Figure 1.**
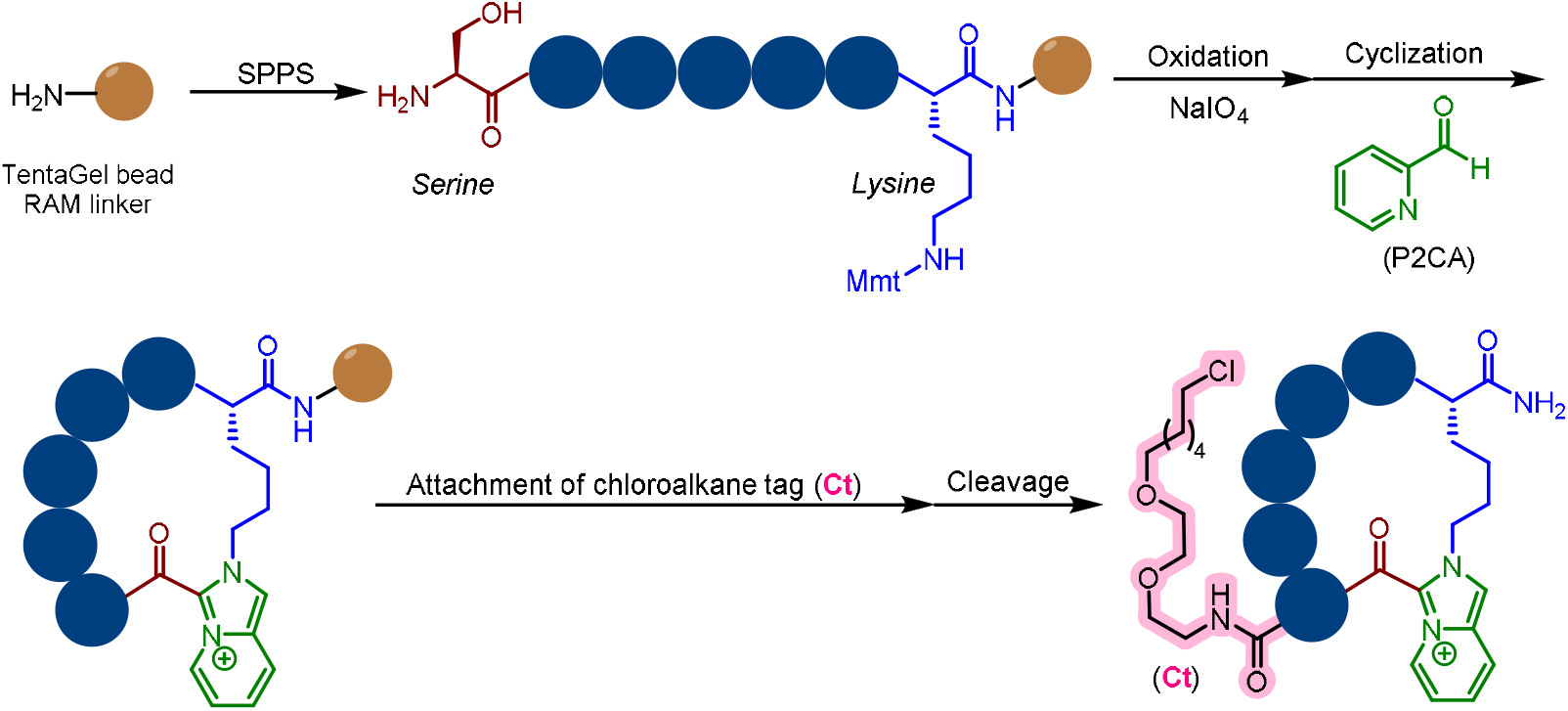
Synthesis of IP^+^-linked MPs bearing a chloroalkane tag (Ct). **Mmt**: Monomethoxytrityl; **Oxidation**: NaIO_4_ (5.0 equiv), MeOH/H_2_O (1:4, v/v), r.t., 1h; **Cyclization**: AcOH/TFE (1:2, v/v), P2CA (3.0 equiv), 37°C, 10 h; **Cleavage**: 100%TFA, r.t., 40 min. The dark circles represent amino acid-derived residues.

### Cell Penetration of IP^+^ MPs Evaluated via CAPA

As mentioned above, one of our motivations in developing IP^+^ as a permeability stimulator was to expand the chemical space of amino acid side chains that could be employed beyond simple hydrophobic residues whilst retaining reasonable cell permeability. Therefore, in the experiments described below, we synthesized IP^+^ MPs containing a broader selection of amino acids, including those with polar side chains, and tested their permeability by CAPA. As a benchmark, the small molecule conjugate **JQ-1**-Ct (**Figure 2A**) was used as a positive control due to its well-characterized cell permeability and validated performance in CAPA.^12^

**Figure 2.**
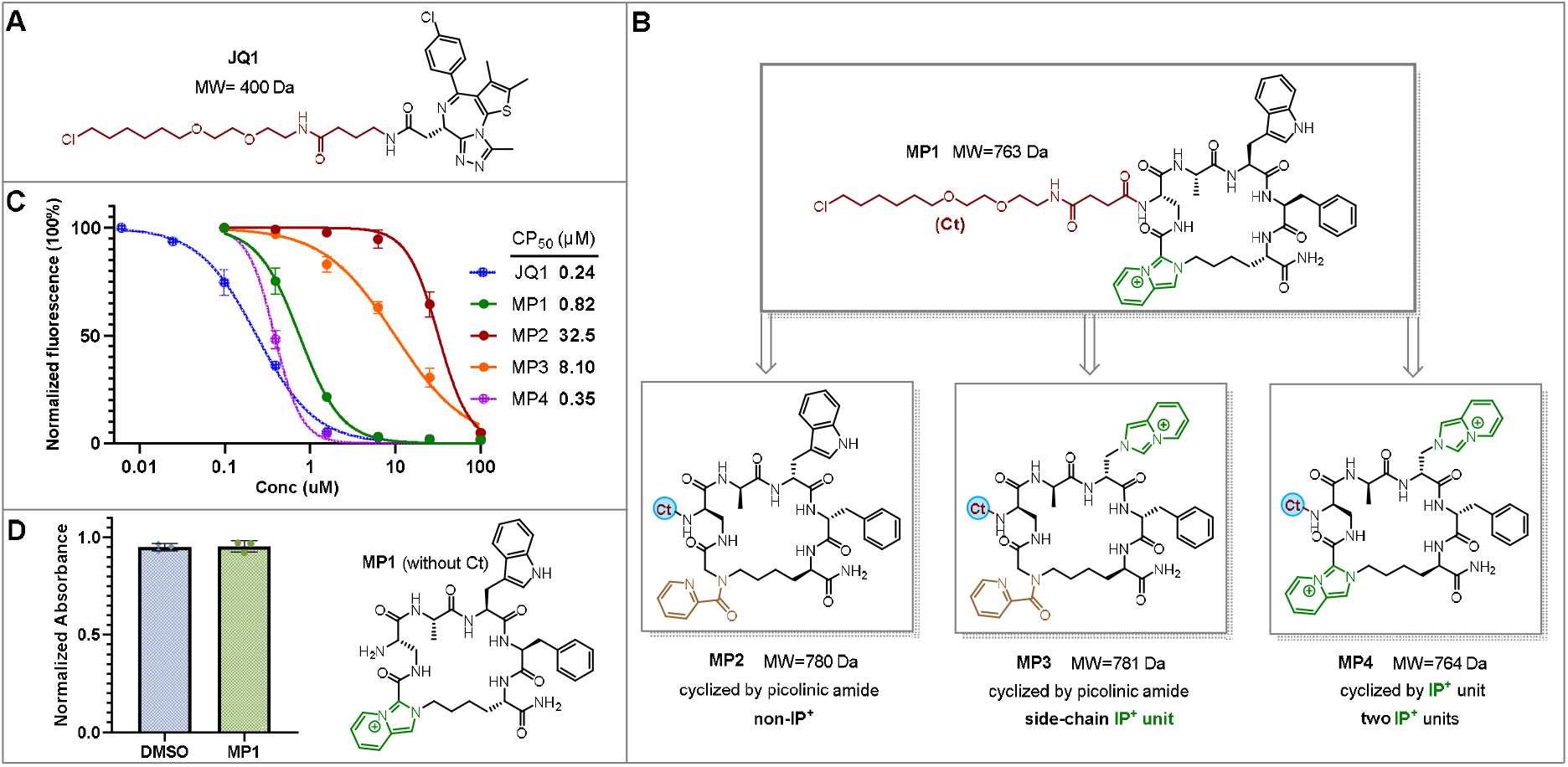
Comparison of the cell permeability of MPs containing one or two IP+ units with that of a drug-like small molecule (JQ1) and a MP lacking the IP+ moiety. **A)** Structure of the Ct conjugate of JQ1. **B)** Structures of the Ct conjugates of the MPs used in this experiment. **C)** CAPA data for the compounds indicated. The results shown are from two independent experiments. CP_50_ curves were calculated by GraphPam Prism using a inhibitor vs. normalized response model. All molecular weights listed in this figure do not include the mass of the Ct. D) Effect of MP1 on the viability of HEK293 cells. Cells were treated with either DMSO (100 μM) or MP1 (100 µM) for 12 hours. Viability was measured by CellTiter Glo 2.0. The data shown in C and D are the results of three independent experiments. Note that in this figure and throughput the remainder of the paper, the molecular weights indicated do not include the mass of the Ct.

A model five-residue, IP^+^-containing, medium-sized macrocyclic peptide-Ct conjugate, **MP1**-Ct (Dap-Ala-Trp-Phe-Lys-NH_2_; MW = 763 Da without the Ct), (**Figure 2B**) was prepared via the solid-phase route outlined in **Figure 1**. This peptide contains a Trp residue, a heterocyclic amino acid that, along with tyrosine,^13^ often makes critical contacts with target proteins^14^ but is usually excluded in designing Cyclosporine-like MPs. As a control, we also created **MP2**-Ct, featuring a picolinic amide linkage in place of the IP^+^ unit. This is the equivalent of opening the imidazole ring and removing the positive charge of the IP^+^ heterocycle. CAPA revealed that **MP1**-Ct exhibited a CP_50_ of 0.82 μM (**Figure 2C**), a value only 3.4-fold poorer than that of **JQ-1**-Ct (CP_50_ = 0.24 μM, **Figure 2C**). In contrast, **MP2**-Ct was found to have a CP_50_ of 32.5 μM, a 40-fold increase compared to **MP1**-Ct. In the course of carrying out these experiments, we observed no indication that any of the MPs tested were toxic to the cells. This was demonstrated directly using a commercial CellTiterGlo assay. Even at a concentration of 100 µM, MP1 had no effect on the viability of HEK293 cells (**Figure 2D**).

We next asked whether the IP^+^ unit would be as effective in promoting cytosolic exposure when positioned on a side chain rather than in the macrocyclic ring. This led to the synthesis of **MP3**-Ct (**Figure 2B**), in which the side chain of a Trp residue was replaced with an IP^+^ moiety, while ring closure was via the picolinic amide linkage (see SI for synthesis details). CAPA showed that **MP3**-Ct (CP_50_ = 8.1 μM) was almost ten-fold less permeable than **MP1**-Ct, indicating that placement of the IP^+^ unit in the ring confers greater cell permeability than locating it on the periphery. Nevertheless, **MP3**-Ct remained more permeable than **MP2**-Ct, with a 4-fold lower CP_50_ value.

We also created **MP4**-Ct, bearing IP+ units at both the ring junction and in a side chain (**Figure 2B**). The CP_50_ of this molecule (0.35 μM) was superior to that of even **MP1**-Ct and nearly equivalent to that of the **JQ-1**-Ct conjugate. The difference between the CP_50_s of **MP4**-Ct and **MP2**-Ct was more than 90-fold.

We next expanded our study to additional IP^+^ MPs with diverse structural features. **MP5**-Ct represents a relatively small macrocycle that would adhere to one of the “Cyclosporine rules” in that it contains only hydrophobic side chains. **MP5**-Ct (CP_50_ = 5.7) was found to be more permeable than its non-IP^+^ counterpart **MP6**-Ct (CP_50_ = 24.5 μM), providing another example of the stimulatory effect of the IP^+^ unit. Interestingly, however, **MP5**-Ct (CP_50_ = 5.7) scored as markedly less permeable than **MP1**-Ct (CP_50_ = 0.82), even given its greater hydrophobicity and lower MW. **MP1** and **MP5** were constructed using different amino acids at the ring junction (Lys and Dap, respectively). To better investigate whether this change in the local environment of the IP^+^ unit might be the cause of this difference, we created **MP7**-Ct and measured its permeability by CAPA. **MP7**-Ct and **MP5**-Ct differ only in the placement of either Lys or Dap at the ring junction. As shown in **Figure 3A, MP7**-Ct was almost four-fold more permeable than **MP5**-Ct, suggesting that placement of the more hydrophobic four carbon chain of lysine at the ring junction is superior to that of the one carbon side chain of Dap in promoting cell permeability.

**Figure 3.**
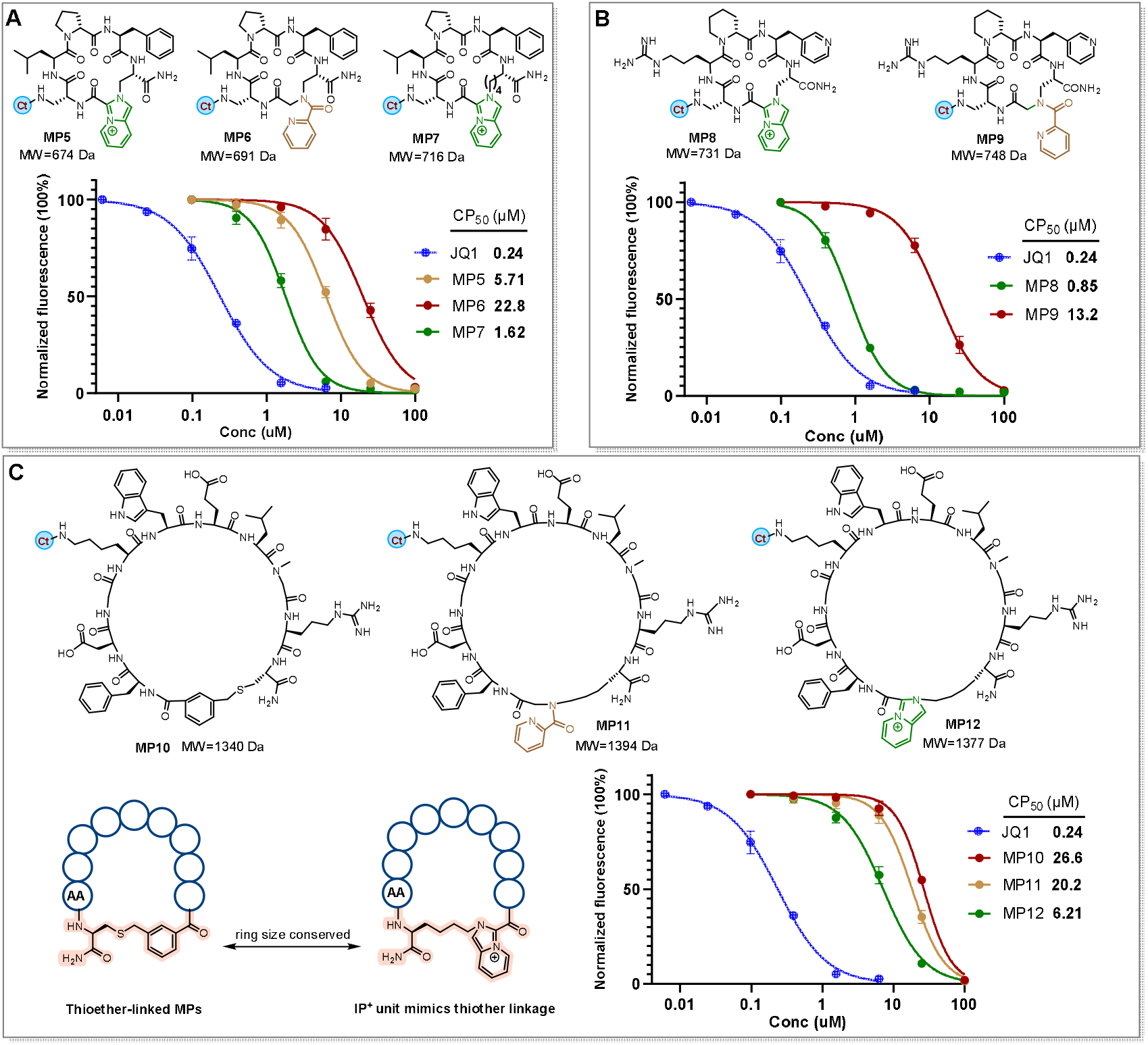
Assessment of the cell permeability of MPs of various sizes and sequence composition. **A**. CAPA data for the Ct conjugates of IP^+^ MPs with solely hydrophobic side chains and derived from molecules containing C-terminal Dap (**MP5**) or Lys (**MP7**) residues, as well as a control MP lacking the IP^+^ unit (**MP6**). **B**. Comparison of the cell permeability of the Ct conjugates of MPs containing polar residues with (**MP8**) and without (**MP9**) an IP^+^ unit in the ring. C. Comparison of the cell permeability of the Ct conjugates of larger MPs containing polar residues that do (**MP12**) or do not (**MP10** and **MP11**) contain an IP^+^ unit. The illustration at the bottom left highlights that the IP^+^ and thioether linkages employed conserve the ring size in these analogues. In all of these experiments, the CAPA data shown are the result of two independent experiments.

We next explored how polar functional groups would influence uptake. We first incorporated arginine (Arg) and 3-pyridylalanine (3-Pal) into a new five-residue macrocycle, **MP8** (**Figure 3B**), which exhibited excellent cell permeability (CP_50_ = 850 nM). Its non-IP^+^ counterpart, **MP9**, displayed dramatically lower permeability, with a CP_50_ over 15-fold higher (13.2 μM). These data demonstrate that IP^+^ macrocycles in this size range can tolerate the inclusion of polar side chains without compromising cell permeability.

### Effect of IP^+^ Incorporation Into Larger MPs

Protein-binding peptides that arise from many phage or mRNA display screens occupy a size range similar to Cyclosporine, which is an 11-residue macrocyclic peptide (MW 1204 Da). ^15^ Therefore, it is of interest to compare the permeability of IP^+^-containing MPs with these characteristics to analogues that lack this unit.

Towards this end, the three MP-Ct conjugates shown in **Figure 3C** were synthesized. All have MWs in excess of 1300 Da and contain several polar side chains, including two carboxylic acid side chains. **MP10**-Ct (**Figure 3C**) was prepared via a Cys-mediated S_N_2 cyclization, forming a benzoyl thioether. IP+-containing **MP12**-Ct and its non-IP+ analogues **MP11**-Ct were made as described above. Not surprisingly, CAPA data (**Figure 3C**) revealed that these large, polar macrocycles were considerably less cell permeable than JQ-1-Ct. Nonetheless, the CP_50_ of **MP12**-Ct was four-fold better than that of **MP10**-Ct and more than three-fold superior to that of **MP11**-Ct indicating that that the stimulatory effect of the IP^+^ unit extends to macrocycles of this size range.

### Elaboration of the IP+ unit impairs cell permeability

Many 2-formyl pyridines with additional substitution are commercially available. Thus, it is straightforward to produce IP^+^ MPs with various additional substitution on the heterocycle.^6^ Since these substitutions could influence the physical and electronic properties of the IP^+^ ring, which, in turn, could alter its interactions with the lipid membrane, it is of interest to examine the effect of IP^+^ ring modifications on the permeability of IP^+^ MPs.

**MP13**-Ct, containing, a “plain”, unsubstituted IP^+^ heterocycle served as the parent compound for this study. **MP14**-Ct was created as a non-IP^+^ control. As expected, CAPA showed that the CP50 of MP13-Ct was much superior to that of MP-14Ct (6.1 vs. 67.6). A series of **MP13** analogues with various substituents, including electron-donating and withdrawing units as well as more hydrophobic groups, on the IP^+^ ring (**Figure 4A**) was constructed using the appropriate substituted 2-formyl pyridine for the ring-closing reaction. Each compound was tested using the CAPA at a single concentration (10 µM), which is close to the CP_50_ of **MP13**-Ct. The results are shown in **Figure 4C**. None of the substitutions improved the permeability of the MP relative to the parent molecule. In fact, all displayed lower permeability, though in most cases the effect was modest, with the chloro-substituted **MP17**-Ct being a notable exception. These data show that, at least within the series of compounds analyzed, that cell permeability cannot be enhanced by elaboration of the IP^+^ ring. However, if one wishes to do so, these data suggest the “permeability penalty” is likely to be modest.

**Figure 4.**
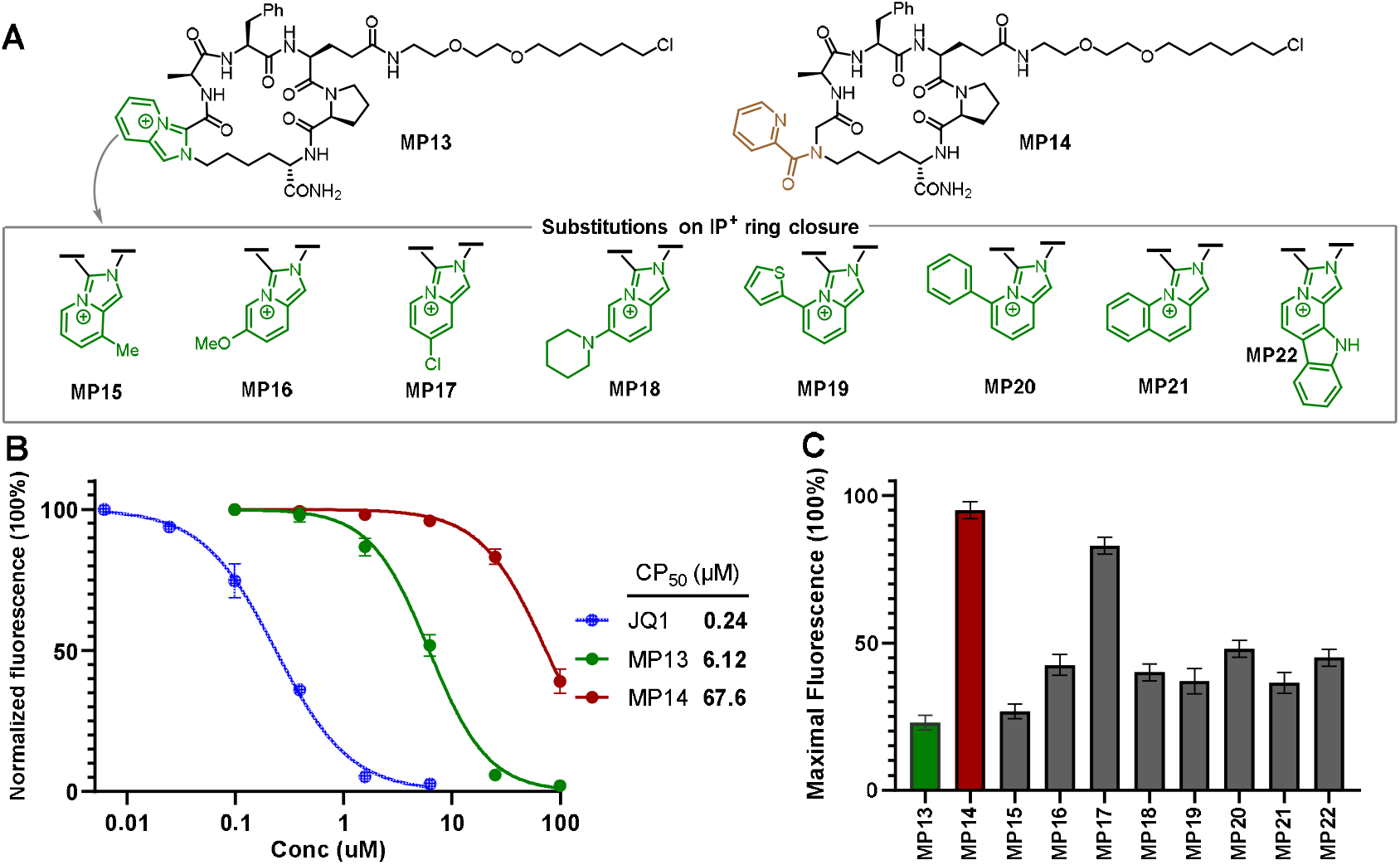
Effect of IP^+^ substitution on the cell permeability of an MP. **A)** Structure of **MP13**-Ct, the non-IP^+^-containing control **MP14**-Ct as well as the various elaborated IP^+^ units that were investigated. **B)** Comparison of the cell permeability of **MP13**-Ct and **MP14**-Ct. The results shown are from two independent experiments. **C)** Single-concentration test of IP^+^ substituted MPs.

### Improving the permeability of a MP inhibitor of the p53-MDM2 interaction

The data above show that in every case tested, inclusion of an IP^+^ unit into an MP improves its cell permeability, in some cases dramatically. This raises the possibility that the cellular activity of existing protein-binding MPs with poor plasma membrane permeability could be improved by incorporation of an IP^+^ ring.

To test this idea, we selected **UNP-6457** (**Figure 5A**), a molecule derived from a screen of a DNA-encoded library of macrocyclic peptides, which potently inhibits the MDM2–p53 interaction *in vitro* (IC_50_ = 8.9 nM) but cannot penetrate cell membranes, resulting in no measurable cellular activity.^8^ Initially, we considered replacing the triazole, the formation of which was used to close the ring, with an IP^+^ unit, but this would alter the macrocycle ring size and potentially disrupt binding. Analysis of the co-crystal structure of **UNP-6457** bound to MDM2 revealed that 3-trifluoromethylphenylalanine, L-Phe, and the benzoyl group derived from amino-proline occupy the central binding pocket, forming key hydrophobic interactions critical for mimicking the α-helical MDM2-binding region of p53. In contrast, the Trp residue is solvent-exposed and does not directly contribute to MDM2 engagement, making it an attractive site for modification. Thus, we generated **MP23** (**Figure 5A**) and its effect on the viability of MCF-7 breast cancer cells was assessed. Loss of the p53-Mdm2 interaction is toxic to these cells.^16^ As shown in Figure 5C, as expected, **UNP-6457** had no effect on MCF-7 viability. In contrast, **MP23** did display measurable cytotoxicity (IC_50_ ≈ 35 µM), indicating a significant improvement in the cell permeability of the macrocycle as a result of the Trp to IP^+^ substitution. Navtemadlin, a clinically tested, orally bioavailable p53–MDM2 inhibitor, which was used as a positive control, was about ten-fold more potent than **MP23**. A linear analogue of **MP23** (**MP23-L**), which would be expected to have negligible affinity for MDM2, had no effect on cell viability.

**Figure 5.**
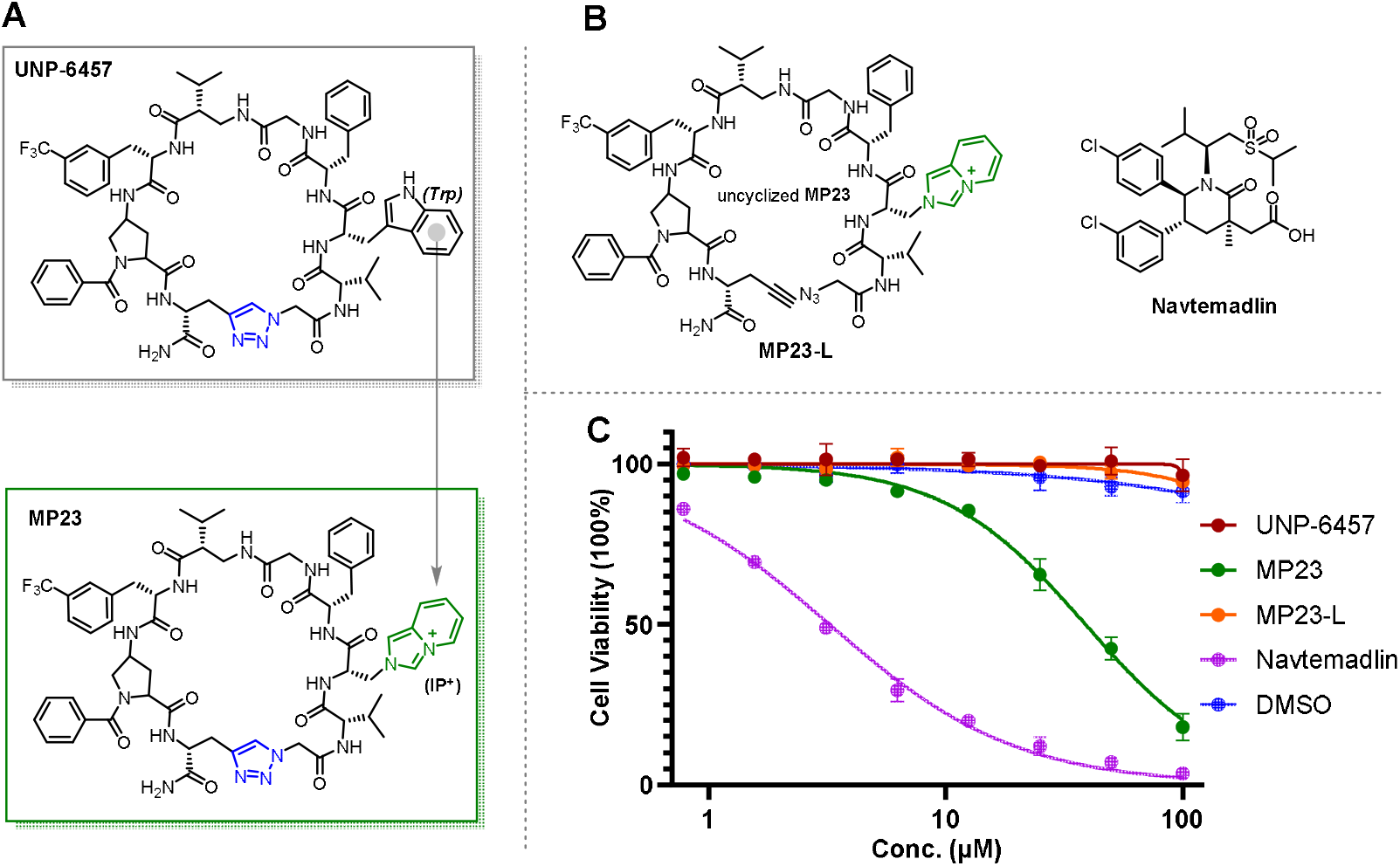
A) Structure of UNP-6457 and MP23. B) Structure of MP23-L and Navtemadlin. C) Cell viability assay for MCF-7. The cells were treated with the testing compounds for 48h.The cell viability assay was measured by CellTiter Glo 2.0 from Promega. Results were reproduced in two independent experiments.

### Mechanistic study of IP^+^ MPs Entry into Cells

The data presented above establish the significant effect of the IP^+^ unit in increasing the cytoplasmic exposure of MPs containing this unit. An important question is whether this occurs by simple diffusion through the plasma membrane or by some type of active transport mechanism such as endocytosis, as is the case for many types of cell-penetrating peptides and related molecules.^17^ In our previous study, many IP^+^ MPs displayed excellent permeability in the PAMPA, a cell-free system devoid of active transport machinery. Nonetheless, we deemed it important to examine this issue in more detail in living cells.

A549 were cells treated with a fluorescently labeled IP^+^ MP, **MP1**-Bodipy (**Figure 6A**). This was done in such a way that an image could be recorded within one minute of exposure (see details in the SI). This revealed an immediate, uniform staining of the plasma membrane with **MP1**-Bodipy (**Figure 6B**). A diffuse cytosolic signal was also detectable at this early time point (**Figure 6B**). Quantitative fluorescence analysis, calibrated against a standard curve, revealed that intracellular **MP1-Bodipy** reached near-equilibrium levels rapidly (**Figure 6C**). To link these imaging results with a functional readout, we used **MP1**-Ct in a time-resolved CAPA experiment. As shown in **Figure 6D, MP1**-Ct alkylation of HaloTag protein was relatively rapid, with kinetics only slightly slower than that seen for **JQ-1**-Ct.

**Figure 6:**
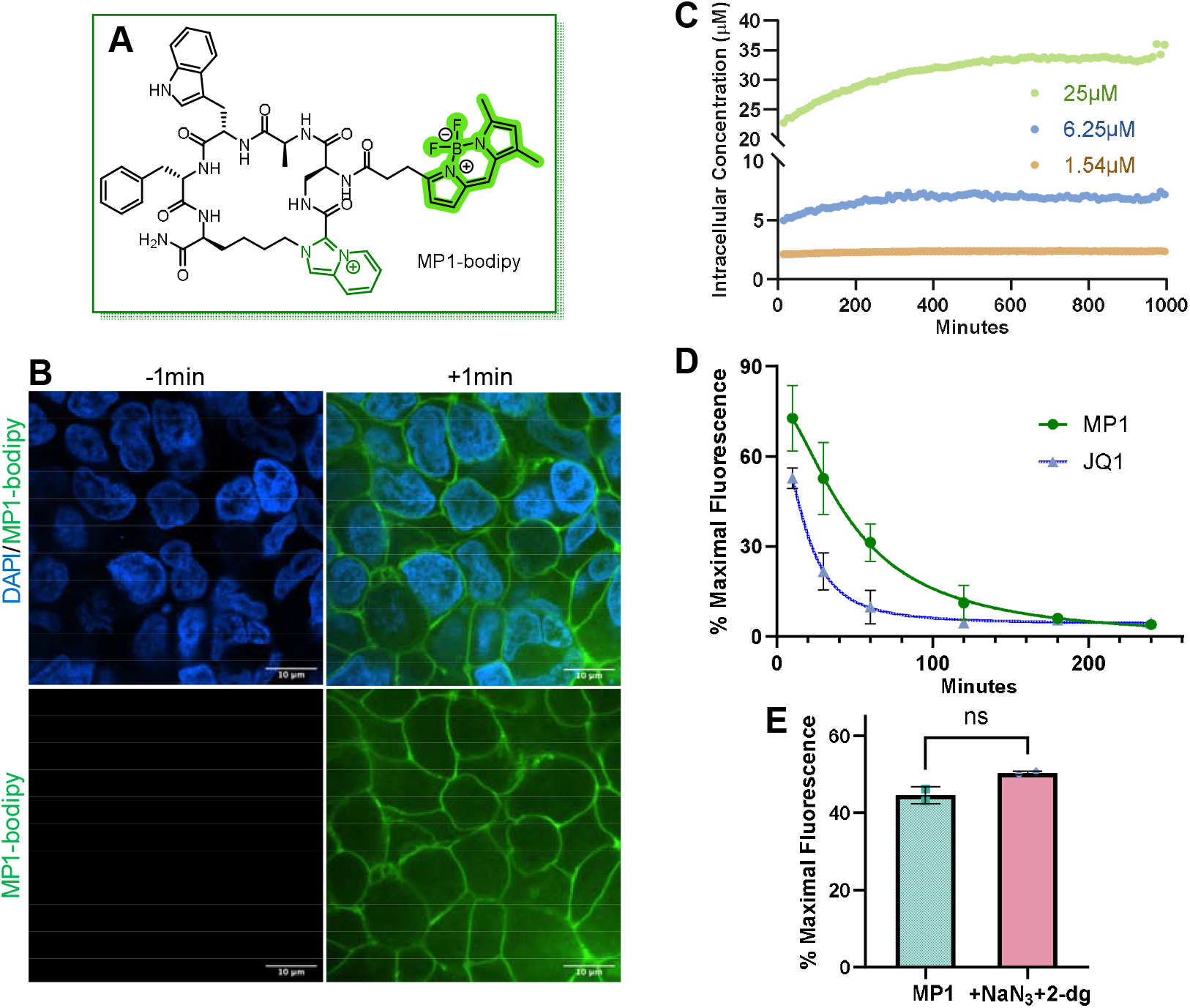
IP^+^ MPs Display Small Molecule Entry Patterns. A) Structure of 1a-bodipy. B) A549 cells 1 min prior to and after 5μM 1a-bodipy treatment. C) HEK293 cell intracellular 1a-bodipy concentration over time. A standard curve was generated by measuring the fluorescence of media at a gradient of 1a-bodipy concentrations. This curve was used to assess the intracellular concentration of 1a-bodipy across time. D) Time-resolved CAPA for MP1 and JQ1. 5μM MP1 and 1.62μM JQ1 were used for this assay. Results were reproduced in two independent experiments. E) CAPA readout in ATP depleting conditions for MP1. Halo-tag expressing cells were treated for 4 hours with 0.77 μM MP1 in the presence or absence of 10mM NaN_3_ and 2-dg. Cellular uptake was assessed by chasing with the same fluorescently-labeled Ct containing molecule used elsewhere in this study. Results were reproduced in two independent experiments.

These two different indicators of rapid entry of the **MP1** derivatives into the cytoplasm would seem to be more compatible with a passive permeability mechanism than active transport. But to address this more directly, cells were treated with **MP1-**bodipy in the presence of the ATP-depleting agents sodium azide (NaN_3_) and 2-deoxyglucose (2-dg), which block active transport-dependent pathways of cellular entry. Quantification of the level of intracellular fluorescence showed that these manipulations had no effect (**Figure 6E**), arguing against an active mechanism of IP^+^ MP entry. We further evaluated potential membrane destabilization or pore formation, classical hallmarks of cationic CPP uptake, by co-treating cells with propidium iodide (PI), a cell impermeable DNA stain. No increase in PI-dependent fluorescence signal was observed for **MP1**-treated cells, while detergent-permeabilized cells displayed robust nuclear staining (see **Supplementary Scheme 15**).

Taken together, these results indicate that IP^+^ MPs associate strongly with the plasma membrane without inducing membrane damage or pore formation and enter cells independently of ATP-dependent mechanisms. We propose that the primary uptake pathway for IP^+^ MPs is passive diffusion, likely facilitated by the combination of the cationic IP^+^ unit and the hydrophobic character of the macrocycle.

### IP^+^ MPs Do Not Co-localize with Mitochondria or Endosomes

Another important question is whether IP^+^ MPs localize to mitochondrial membranes, as is the case for many other positively charged compounds.^18^ To investigate this, HEK293 cells were treated with **MP1**-bodipy and stained with MitoTracker DeepRed, a well-established mitochondrial marker. High-resolution confocal imaging revealed no significant colocalization between the MP1-bodipyand mitochondrial signals as quantified by the widely used ImageJ Coloc 2 plugin.(**Figure 7A&B**). Instead, fluorescence was broadly distributed throughout the cytoplasm.

**Figure 7:**
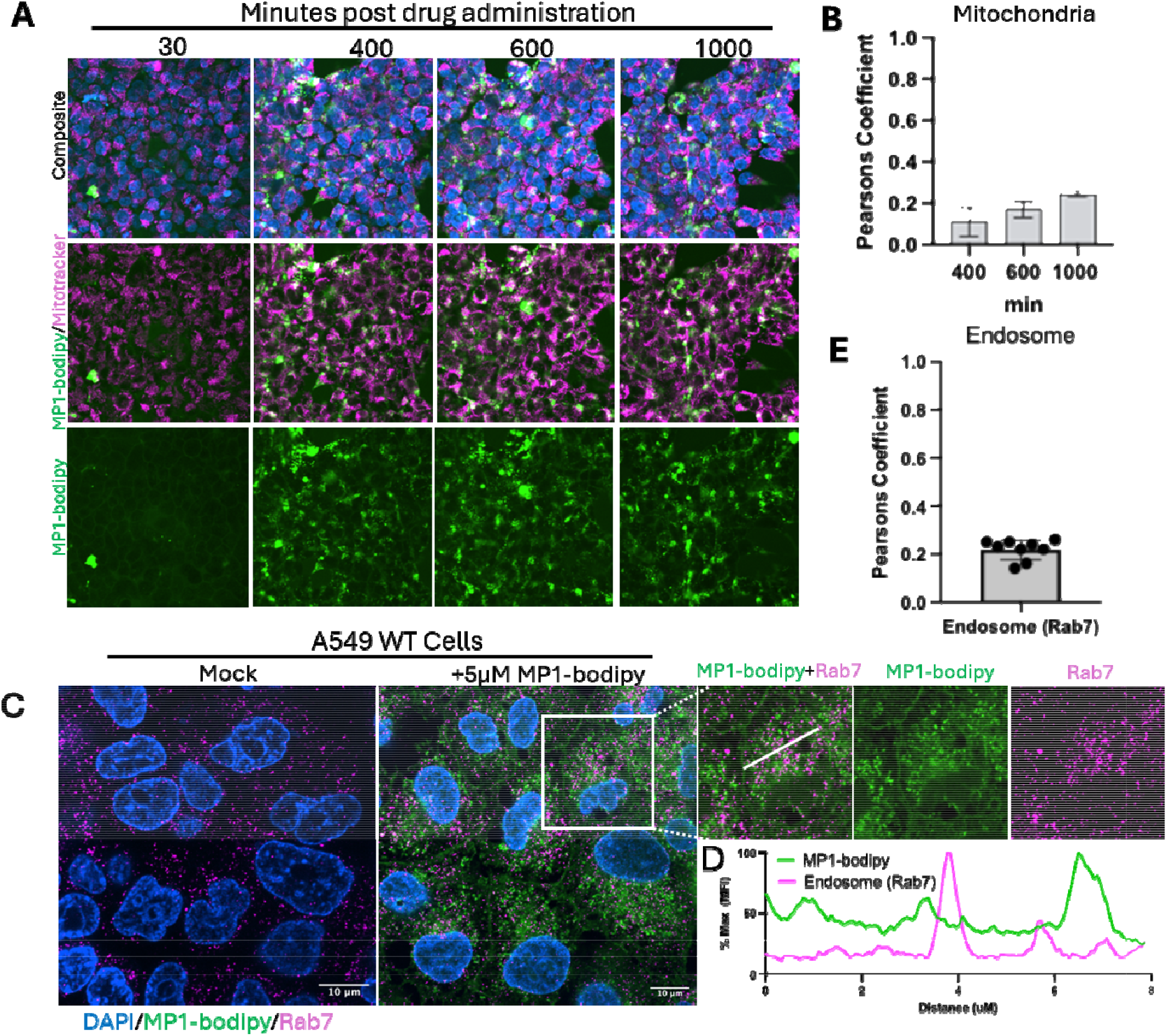
IP^+^ MPs Do Not Colocalize with Mitochondria or Endosomes. (A) Live HEK293 cells imaging following treatment with treatment with **MP1**-bodipy (6.25 µM) in the presence of MitoTracker Deep Red. (B) Quantification of **MP1-bodipy** and MitoTracker colocalization. Values closer to 1 indicate a high degree of colocalization. (C) Determination of the degree of co-localizaiotn of **MP1**-Bodipy with Rab7, a marker for late endosomes. Cells were exposed to **MP1**-bodipy (6.25 µM) for four hours, then fixed, permeabilized, and stained with an anti-Rab7 antibody and fluorescently labeled secondary antibody. D. Colocalization was evaluated by a line trace representing MFI of the two signals across the diameter of a single cell.

While earlier experiments argued against endocytic uptake of IP^+^ MPs, we also examined possible colocalization of **MP1**-bodipy in A549 cells with Rab7, a marker for late endosomes.^19^ Briefly, cells were incubated with **MP1**-bodipy for four hours, fixed, then stained with an anti-Rab7 antibody and a fluorescently labeled secondary antibody. Confocal imaging revealed no significant colocalization between the fluorescent signal of **MP1**-bodipy and that of the labeled anti-Rab7 antibody.(**Figure 7C & 7D**).

## Discussion

MPs are of increasing interest in the pharmaceutical industry, particularly for engaging difficult protein targets. However, an enduring challenge in the field is to improve the generally poor bioavailability and cell permeability of these molecules.^20^ As described above, most researchers in the field have focused on strategies that eliminate as many N-H amide bonds as possible through methylation or the formation of intramolecular hydrogen bonds, and generally increasing the lipophilicity of the molecule. ^2, 21^ Recently, we explored a different strategy. Rather than removing the impediments to passage across the membrane, we sought to identify moieties that would intrinsically increase the passive membrane permeability of MPs. Towards this end we showed that formation of an IP^+^ heterocycle is a highly efficient and general method for the key ring closure step in the synthesis of MPs.^6^ While several other efficient methods for closing the ring of MPs exist,^22^ we postulated that the permanent positive charge of the IP^+^ would lead the MPs to concentrate at the surface of negatively charged phospholipid membrane and then the hydrophobic nature of the heterocycle would allow the compound to more efficiently slip through the bilayer. We were gratified to find that in a group of twenty randomly chosen IP^+^ MPs with MWs ranging from 650 Da to 1300 Da the passive membrane permeability of about half of the molecules scored as drug-like, as determined by PAMPA, and another 25% displayed moderate permeability. Analogues lacking an IP^+^ unit traversed the membrane 10-to 100-fold more slowly.^6^

This study focused on evaluating the efficiency and mechanism of entry of IP^+^ MPs into living cells. As predicted by the PAMPA data, we find that in every case examined, IP^+^ MPs are more cell permeable than analogues lacking this unit, as determined by the CAPA. In some cases, the difference is dramatic. For example, as shown in Figure 2, IP^+^-containing **MP1**-Ct is approximately 40 times more permeable than its picolinic acid-containing analogue **MP2**-Ct. The inclusion of an additional IP^+^ unit into **MP1**, by replacing the indole ring of the Trp with an IP^+^ unit, increased the permeability of the molecule another two-fold. **MP4**-Ct, with a MW of 764 Da (excepting the Ct), was almost as cell permeable as the Ct derivative of **JQ1**, a typical drug-like small molecule. These experiments also suggested that the IP^+^ unit is more effective when integrated into the macrocycle than as a side chain unit, though more data points will be necessary to determine this definitively.

Larger or more polar macrocycles also benefitted from IP^+^ incorporation (see Figures 3 and 5), suggesting the approach is broadly applicable across diverse peptide scaffolds. Indeed, we created an IP^+^-containing analogue of **UNP6457**, a nine-residue MP that potently antagonizes binding of p53 to Mdm2 *in vitro* but has no cellular activity due to its inability to cross membranes.^8^ This molecule (**MP23**) displayed readily detectable cytotoxicity to MCF-7 breast cancer cells (**Figure 5**), indicating a significant IP^+^-dependent increase in permeability.

However, it is also important to note that the data in **Figures 3** and **5** make clear that the IP^+^ unit is not a “magic wand” that will transform any MP into a molecule with small molecule-like permeability. For example, the cytotoxic IC_50_ of **MP23** remains far higher than the biochemical potency of UNP6457. While this is not an “apples vs. apples” comparison, it suggests that even for **MP23**, entry of the molecule into cells remains a limiting factor. In addition, while **MP12**-Ct, a larger molecule with a MW of almost 1400 (again excepting the Ct) demonstrated better permeability than it’s picolinic acid analogue in the CAPA, the CP_50_ value for **MP12**-Ct (6.2 µM) was about 25-fold higher than that of **JQ-1**-Ct (0.24 µM). Having said that, it is still remarkable that **MP12**-Ct displays a CP_50_ in the low µM range given its high MW and the fact it contains Glu, Asp, Trp and Arg residues (**Figure 3**).

The imaging experiments and CAPA data shown in **Figure 6** are consistent with an almost immediate concentration of the IP^+^ MP on the plasma membrane (**Figure 6B**), followed by rapid diffusion into the cell. Passive diffusion, rather than an active transport mechanism such as endocytosis is supported by several lines of evidence. These include the rapid kinetics by which the intracellular concentration of the molecule rises to a level similar to that in the extracellular space (Figure 6C), the insensitivity of IP^+^ MP cell entry to ATP poisons (Figure 6E), and the lack of co-localization of the IP^+^ MP with Rab7, an endosome marker (**Figure 7C**). Also, as mentioned above, many IP^+^ MPs score as highly membrane permeable in PAMPA^6^ where there is no possibility of active transport.

Importantly, once **MP1**-Ct enters cells, it does not localize to mitochondrial membranes (**Figure 7A and B**), as is the case for many other positively charged molecules, especially those that are hydrophobic.^18^ This is consistent with the CAPA data, which score molecules able to react with cytoplasmic Halotag protein. However, inspection of micrographs such as those shown in Figure 7, do show concentration of **MP1**-Bodipy into puncta at later times, in addition to diffuse staining throughput the cell. At present we do not know the mechanistic basis of this phenomenon, other than the puncta do not include mitochondria or endosomes. Addressing this point will the focus of future studies. It is important to point out however, that in all of our experiments we have never observed toxicity when cells are treated with IP^+^ MPs (also see Figure 2). So, these puncta, whatever they are, do not interfere with any critical cellular phenomena. We also cannot rule out the possibility that they are unique to the bodipy-labeled compound.

In summary, we have shown that inclusion of an IP^+^ unit into an MP reliably increases the entry of these Bro5^4^ compounds into cells through passive membrane permeation. We anticipate that this discovery will facilitate the development of MP probe molecules and drug candidates that target intracellular proteins.

## Supporting Information

Description of methods, supplementary figures and data for compound characterization (PDF).

## Author contributions

T.K and B.L. conceptualized this study and experiments. B.L. carried out the design and chemical synthesis. J.P. and B.L. performed and analyzed the CAPA experiments. J.D. and B.L. performed the cell viability assay. S.B. and J.P. carried out and analyzed the imaging experiments. B.L. wrote the manuscript. T.K. edited the manuscript and acquired funding for this study. The entire study was supervised by T.K. and J.M.B. All authors contributed to the analysis and interpretation of results.

## References

(1) Johns, D. G.; Campeau, L. C.; Banka, P.; Bautmans, A.; Bueters, T.; Bianchi, E.; Branca, D.; Bulger, P. G.; Crevecoeur, I.; Ding, F. X.; et al. Orally Bioavailable Macrocyclic Peptide That Inhibits Binding of PCSK9 to the Low Density Lipoprotein Receptor. Circulation 2023, 148 (2), 144-158. Iskandar, S. E.; Bowers, A. A. mRNA Display Reaches for the Clinic with New PCSK9 Inhibitor. ACS Medicinal Chemistry Letters 2022, 13 (9), 1379–1383.

(2) Ohta, A.; Tanada, M.; Shinohara, S.; Morita, Y.; Nakano, K.; Yamagishi, Y.; Takano, R.; Kariyuki, S.; Iida, T.; Matsuo, A.; et al. Validation of a New Methodology to Create Oral Drugs beyond the Rule of 5 for Intracellular Tough Targets. Journal of the American Chemical Society 2023.

(3) Stein Gold, L.; Eyerich, K.; Merola, J. F.; Torres, J.; Coates, L. C.; Allegretti, J. R. Oral Peptide Therapeutics as an Emerging Treatment Modality in Immune-Mediated Inflammatory Diseases: A Narrative Review. Adv Ther 2025, 42 (7), 3158–3172.

(4) Lipinski, C. A. Drug-like properties and the causes of poor solubility and poor permeability. Journal of pharmacological and toxicological methods 2000, 44 (1), 235–249. Lipinski, C. A.; Lombardo, F.; Dominy, B. W.; Feeney, P. J. Experimental and computational approaches to estimate solubility and permeability in drug discovery and developmental settings. Adv. Drug Deliv. Rev. 1997, 23, 3–25.

(5) Hutt, J. T.; Aron, Z. D. Efficient, Single-Step Access to Imidazo[1,5-a]pyridine N-Heterocyclic Carbene Precursors. Organic letters 2011, 13 (19), 5256–5259.

(6) Li, B.; Parker, J.; Tong, J.; Kodadek, T. Synthesis of membrane permeable macrocyclic peptides via imidazopyridinium grafting. J. Amer. Chem. Soc. 2024, 146, 14633–14644.

(7) Ono, T.; Tabata, K. V.; Goto, Y.; Saito, Y.; Suga, H.; Noji, H.; Morimoto, J.; Sando, S. Label-free quantification of passive membrane permeability of cyclic peptides across lipid bilayers: penetration speed of cyclosporin A across lipid bilayers. Chem Sci 2023, 14 (2), 345–349.

(8) Silvestri, A. P.; Zhang, Q.; Ping, Y.; Muir, E. W.; Zhao, J.; Chakka, S. K.; Wang, G.; Bray, W. M.; Chen, W.; Fribourgh, J. L.; et al. DNA-Encoded Macrocyclic Peptide Libraries Enable the Discovery of a Neutral MDM2–p53 Inhibitor. ACS Medicinal Chemistry Letters 2023, 14 (6), 820–826.

(9) Deprey, K.; Kritzer, J. A. Quantitative measurement of cytosolic penetration using the chloroalkane penetration assay. Methods Enzymol 2020, 641, 277–309.

(10) Los, G. V.; Encell, L. P.; McDougall, M. G.; Hartzell, D. D.; Karassina, N.; Zimprich, C.; Wood, M. G.; Learish, R.; Ohana, R. F.; Urh, M.; et al. HaloTag: a novel protein labeling technology for cell imaging and protein analysis. ACS chemical biology 2008, 3 (6), 373–382.

(11) Geoghegan, K. F.; Stroh, J. G. Site-directed conjugation of nonpeptide groups to peptides and proteins via periodate oxidation of 2-amino alcohol. Application to modification of a N-terminal serine. Bioconj. Chem. 1992, 3, 138–146.

(12) Filippakopoulos, P.; Qi, J.; Picaud, S.; Shen, Y.; Smith, W. B.; Fedorov, O.; Morse, E. M.; Keates, T.; Hickman, T. T.; Felletar, I.; et al. Selective inhibition of BET bromodomains. Nature 2010, 468 (7327), 1067-1073. Foley, C. A.; Potjewyd, F.; Lamb, K. N.; James, L. I.; Frye, S. V. Assessing the Cell Permeability of Bivalent Chemical Degraders Using the Chloroalkane Penetration Assay. ACS chemical biology 2020, 15 (1), 290–295.

(13) Fellouse, F. A.; Wiesmann, C.; Sidhu, S. S. Synthetic antibodies from a four-amino-acid code: a dominant role for tyrosine in antigen recognition. Proc Natl Acad Sci U S A 2004, 101 (34), 12467–12472, Research Support, U.S. Gov’t, Non-P.H.S. Research Support, U.S. Gov’t, P.H.S.

(14) Shao, J.; Kuiper, B. P.; Thunnissen, A.-M. W. H.; Cool, R. H.; Zhou, L.; Huang, C.; Dijkstra, B. W.; Broos, J. The Role of Tryptophan in π Interactions in Proteins: An Experimental Approach. Journal of the American Chemical Society 2022, 144 (30), 13815–13822.

(15) Jaroszewicz, W.; Morcinek-Orłowska, J.; Pierzynowska, K.; Gaffke, L.; Węgrzyn, G. Phage display and other peptide display technologies. FEMS Microbiology Reviews 2021, 46 (2). Goto, Y.; Suga, H. The RaPID Platform for the Discovery of Pseudo-Natural Macrocyclic Peptides. Accounts Chem Res 2021, 54 (18), 3604–3617.

(16) Shangary, S.; Wang, S. Small-molecule inhibitors of the MDM2-p53 protein-protein interaction to reactivate p53 function: a novel approach for cancer therapy. Annu Rev Pharmacol Toxicol 2009, 49, 223–241.

(17) Nagahara, H.; Vocero-Akbani, A. M.; Snyder, E. L.; Ho, A.; Latham, D. G.; Lissy, N. A.; Becker-Hapak, M.; Ezhevsky, S. A.; Dowdy, S. F. Transduction of full-length TAT fusion proteins into mammalian cells: TAT-p27^Kip1^ induces cell migration. Nature Biotechnol. 1998, 4, 1449-1452. Schwarze, S. R.; Ho, A.; Vocero-Akbani, A.; Dowdy, S. F. In vivo protein transduction: delivery of a biologically active protein into the mouse. Science 1999, 285, 1569-1572. Wender, P. A.; Mitchell, D. J.; Pattabiraman, K.; Pelkey, E. T.; Steinman, L.; Rothbard, J. B. The Design, Synthesis, and Evaluation of Molecules That Enable or Enhance Cellular Uptake: Peptoid Molecular Transporters. Proc. Natl. Acad. Sci. USA 2000, 97, 13003–13008. Futaki, S.; Suzuki, T.; Ohashi, W.; Yagami, T.; Tanaka, S.; Ueda, K.; Sugiura, Y. Arginine-rich Peptides: AN ABUNDANT SOURCE OF MEMBRANE-PERMEABLE PEPTIDES HAVING POTENTIAL AS CARRIERS FOR INTRACELLULAR PROTEIN DELIVERY*. Journal of Biological Chemistry 2001, 276 (8), 5836-5840. Qian, Z.; LaRochelle, J. R.; Jiang, B.; Lian, W.; Hard, R. L.; Selner, N. G.; Luechapanichkul, R.; Barrios, A. M.; Pei, D. Early endosomal escape of a cyclic cell-penetrating peptide allows effective cytosolic cargo delivery. Biochemistry 2014, 53 (24), 4034–4046. Kauffman, W. B.; Fuselier, T.; He, J.; Wimley, W. C. Mechanism Matters: A Taxonomy of Cell Penetrating Peptides. Trends Biochem Sci 2015, 40 (12), 749–764.

(18) Wang, H.; Fang, B.; Peng, B.; Wang, L.; Xue, Y.; Bai, H.; Lu, S.; Voelcker, N. H.; Li, L.; Fu, L.; et al. Recent Advances in Chemical Biology of Mitochondria Targeting. Front Chem 2021, 9, 683220. Novgorodov, S. A.; Szulc, Z. M.; Luberto, C.; Jones, J. A.; Bielawski, J.; Bielawska, A.; Hannun, Y. A.; Obeid, L. M. Positively charged ceramide is a potent inducer of mitochondrial permeabilization. J Biol Chem 2005, 280 (16), 16096–16105.

(19) Vanlandingham, P. A.; Ceresa, B. P. Rab7 regulates late endocytic trafficking downstream of multivesicular body biogenesis and cargo sequestration. J Biol Chem 2009, 284 (18), 12110–12124.

(20) Vinogradov, A. A.; Yin, Y.; Suga, H. Macrocyclic Peptides as Drug Candidates: Recent Progress and Remaining Challenges. Journal of the American Chemical Society 2019, 141 (10), 4167-4181. Buckton, L. K.; Rahimi, M. N.; McAlpine, S. R. Frontispiece: Cyclic Peptides as Drugs for Intracellular Targets: The Next Frontier in Peptide Therapeutic Development. Chemistry – A European Journal 2021, 27 (5).

(21) White, T. R.; Renzelman, C. M.; Rand, A. C.; Rezai, T.; McEwen, C. M.; Gelev, V. M.; Turner, R. A.; Linington, R. G.; Leung, S. S.; Kalgutkar, A. S.; et al. On-resin N-methylation of cyclic peptides for discovery of orally bioavailable scaffolds. Nature Chem Biol 2011, 7 (11), 810-817. Boehm, M.; Beaumont, K.; Jones, R.; Kalgutkar, A. S.; Zhang, L.; Atkinson, K.; Bai, G.; Brown, J. A.; Eng, H.; Goetz, G. H.; et al. Discovery of Potent and Orally Bioavailable Macrocyclic Peptide-Peptoid Hybrid CXCR7 Modulators. J Med Chem 2017, 60 (23), 9653-9663. Koesema, E.; Roy, A.; Paciaroni, N. G.; Coito, C.; Tokmina-Roszyk, M.; Kodadek, T. Synthesis and Screening of a DNA-Encoded Library of Non-Peptidic Macrocycles. Angew Chem Int Ed Engl 2022, e202116999. Zhang, P.; Koch, G.; Zhang, Y.; Yang, K.; Lokey, R. S. DNA-Compatible Conditions for the Formation of N-Methyl Peptide Bonds. ACS Omega 2023, 8 (26), 23477-23483. Faris, J. H.; Adaligil, E.; Popovych, N.; Ono, S.; Takahashi, M.; Nguyen, H.; Plise, E.; Taechalertpaisarn, J.; Lee, H.-W.; Koehler, M. F. T.; et al. Membrane Permeability in a Large Macrocyclic Peptide Driven by a Saddle-Shaped Conformation. Journal of the American Chemical Society 2024, 146 (7), 4582–4591.

(22) Yudin, A. K. Macrocycles: lessons from the distant past, recent developments, and future directions. Chem Sci 2015, 6 (1), 30–49, 10.1039/C4SC03089C. Adebomi, V.; Cohen, R. D.; Wills, R.; Chavers, H. A. H.; Martin, G. E.; Raj, M. CyClick Chemistry for the Synthesis of Cyclic Peptides. Angewandte Chemie International Edition 2019, 58 (52), 19073–19080, 10.1002/anie.201911900. Fang, P.; Pang, W.-K.; Xuan, S.; Chan, W.-L.; Leung, K. C.-F. Recent advances in peptide macrocyclization strategies. Chemical Society Reviews 2024, 53 (24), 11725–11771, 10.1039/D3CS01066J.

