## Supplementary Information for "Macrocyclic Peptides Containing an Imidazopyridinium (IP^+^) Unit Display Enhanced Passive Cell Permeability"

#### General Information

Synthetic TentaGel beads were purchased from Rapp Polymere GmbH (Germany). Disposable reaction columns (Intavis AG) were used as reaction vessels for solid phase peptide synthesis. HPLC was carried out on a Waters systems equipped with a Waters 1525 binary HPLC pumps and a 2487 dual λ absorbance etector, or a 2998 photodiode array detector. The mobile phase comprised of buffer A ($H_2O$ containing 0.1% trifluoroacetic acid (TFA)) and buffer B ($CH_3CN$ containing 0.1% TFA). Analytical HPLC was conducted using a Vydac C-18 column (5 µm, 250 × 4.6 mm, Alltech, Deerfield, IL) at a flow rate of 1.0 mL/min with UV detection at 220 and 254 nm. All steps involving water utilized distilled water filtered through a Barnstead Nanopure filtration system (Thermo Scientific). LC-MS analysis was carried out by Agilent 1100 Series equipped with SBC18 column, PDA detector and a linear gradient of 5% acetonitrile to 95% acetonitrile with 0.05% formic acid.

#### General solid phase synthesis procedures

**Solid phase peptide synthesis:** The Fmoc group was deprotected by 20% piperidine in DMF (2 x 10 min). The beads were extensively washed by DCM (2 x 1 min), DMF (2 x 1 min); Fmoc amino acid (5.0 equiv to resin loading), Oxyma (5.0 equiv) and DIC (5.5 eqiv) were mixed in NMP for 5 min. Afterward, the resulting solution was added to the beads and the reaction was shaken for 1 h at 37°C;

**Oxidation:** After deprotection of Fmoc on N-terminal Ser, a solution of $H_2O$ containing $NaIO_4$ (5.0 equiv) was added to the beads, the reaction was shaken for 1 h at room temperature. Once the reaction was done, the beads were successively washed by $H_2O$ (5 x 1 min), 50% MeOH-$H_2O$ (2 x 1 min), MeOH (2 x 1 min), DCM (2 x 1 min);

**Deprotection of Mmt:** After the Ser was oxidized by $NaIO_4$, the beads were treated with a solution of 30% HFIP/DCM (3 x 30 min). Then the beads were extensively washed by DCM (3 x 1 min).

**Cyclization:** Once the Mmt group was totally removed, the pyridine-2-carboxaldehyde and its analogues were dissolved in AcOH/TFE (1:1 v/v) to give a clear solution, which was mixed with the beads. The reaction was shaken at 37 °C for 10 hours. Then the beads were washed by DMF (2 x 1 min), DCM (2 x 1 min) and $Et_2O$ (2 x 1 min).

**Attachment of chloroalkane tag**: For macrocyclic peptide containing an amine, chloroalkane carboxylate was used for attachment via amide coupling; for macrocyclic peptide containing a carboxylic acid, Ct-amine was used for attachment

**Cleavage and purification:** 100%TFA was added to the beads for 40 min. Then the resin was washed again with TFA. The combined cleavage solutions were volatilized under a stream of argon to give the resulting cyclized peptide, which was subsequently dissolved in water/acetonitrile and purified by reverse-phase HPLC.

### Synthesis of MP1

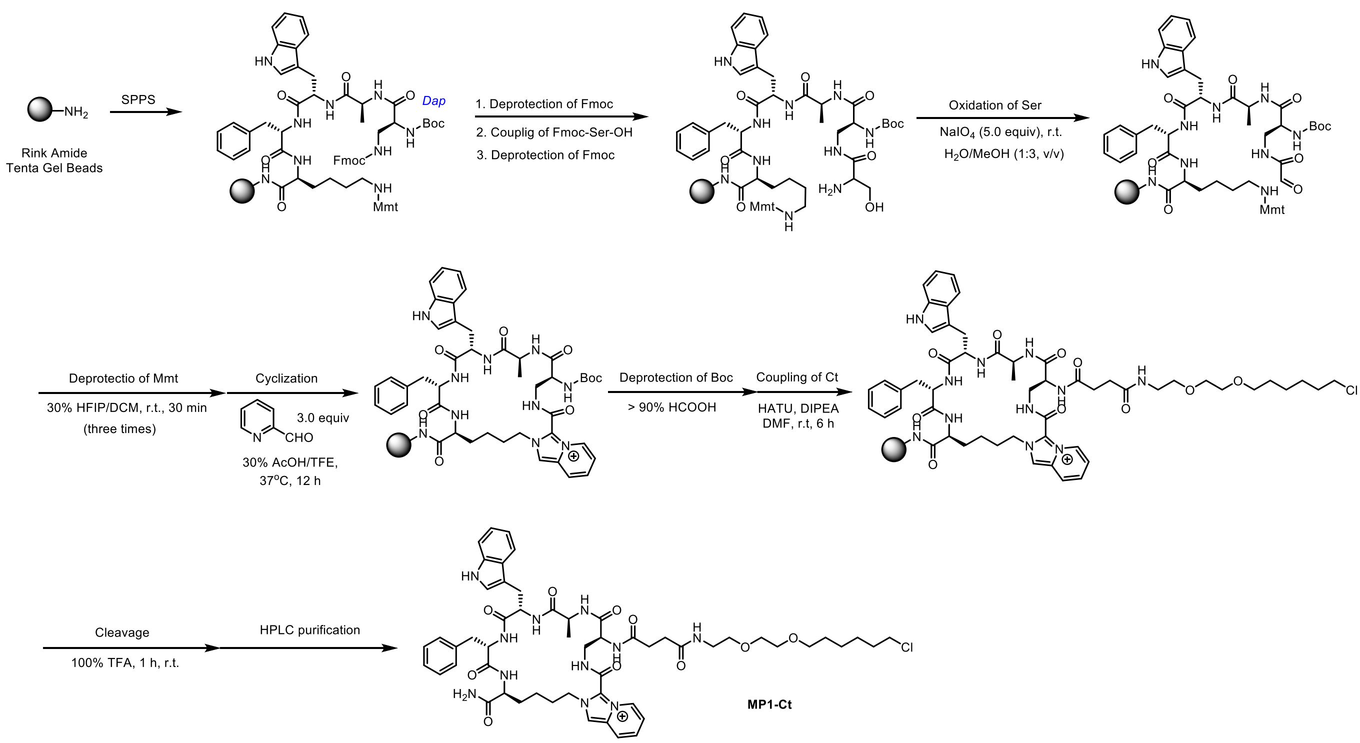


**Supplementary Scheme 1.** Synthetic route to MP1-Ct

Purified HPLC (210 nm) and HRMS of MP1-Ct

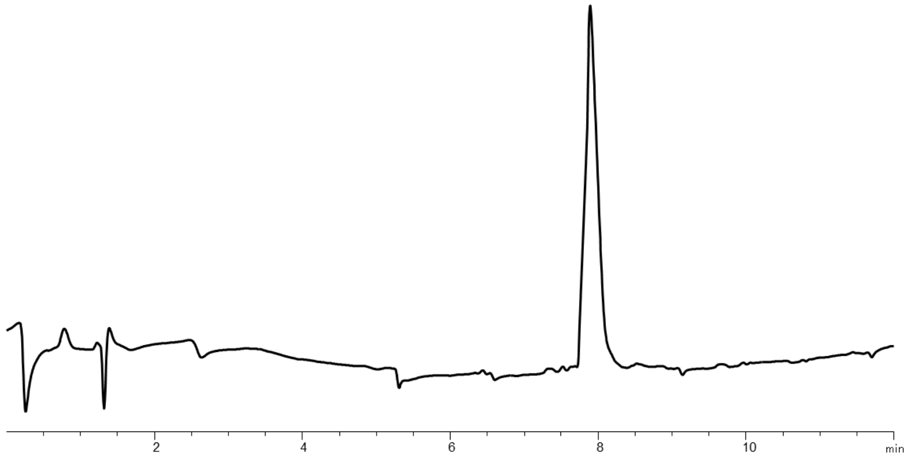


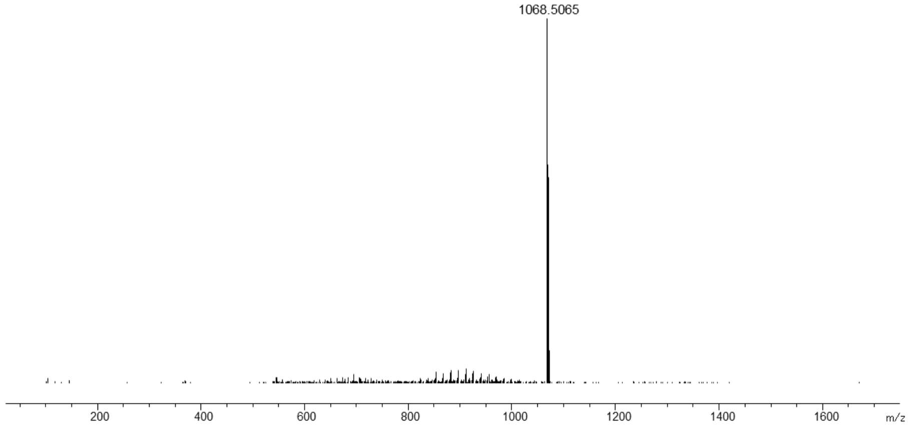

### Synthesis of MP2

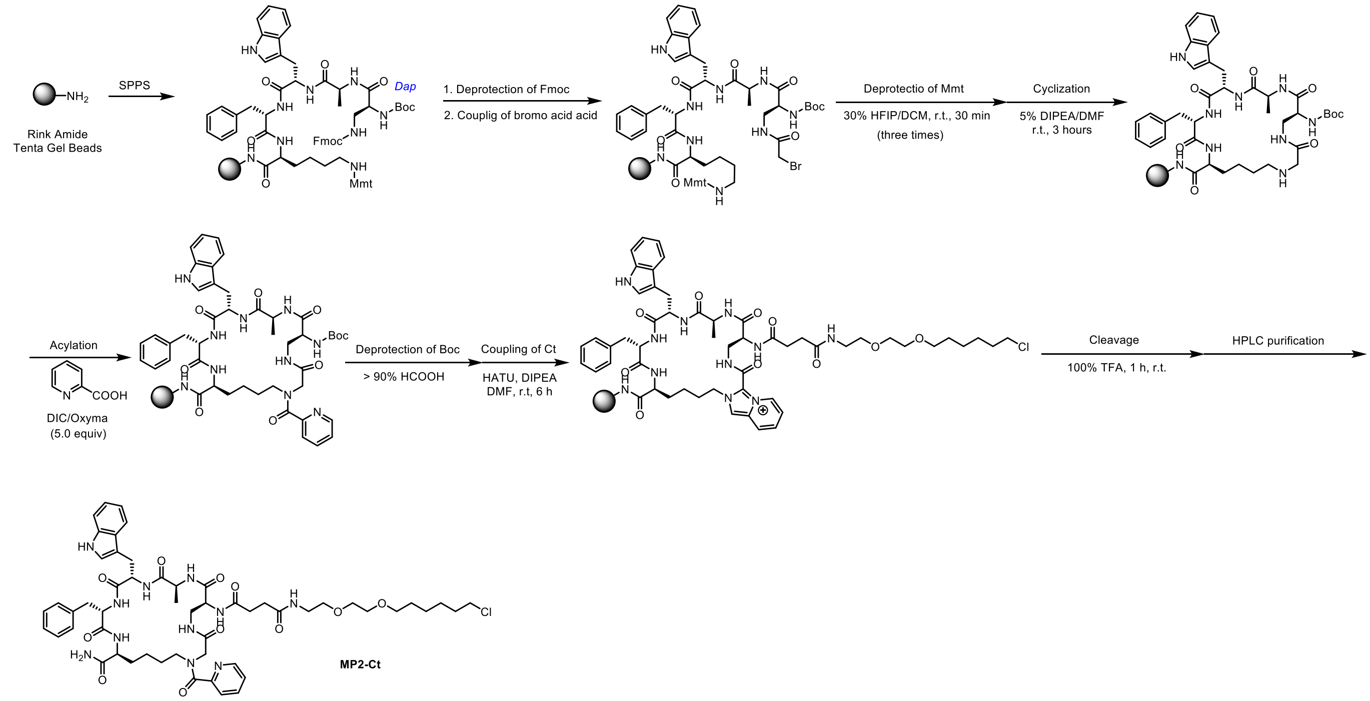


**Supplementary Scheme 2.** Synthetic route to **MP2-Ct**

Purified HPLC (210 nm) and HRMS of MP2-Ct:

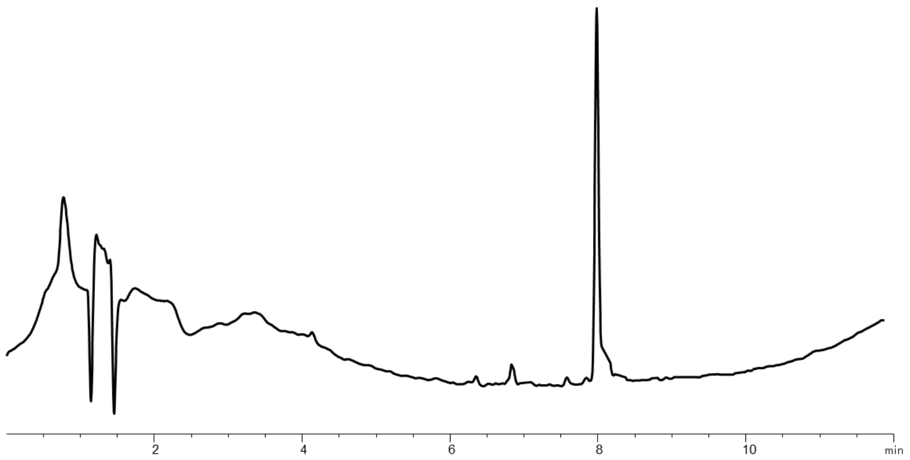


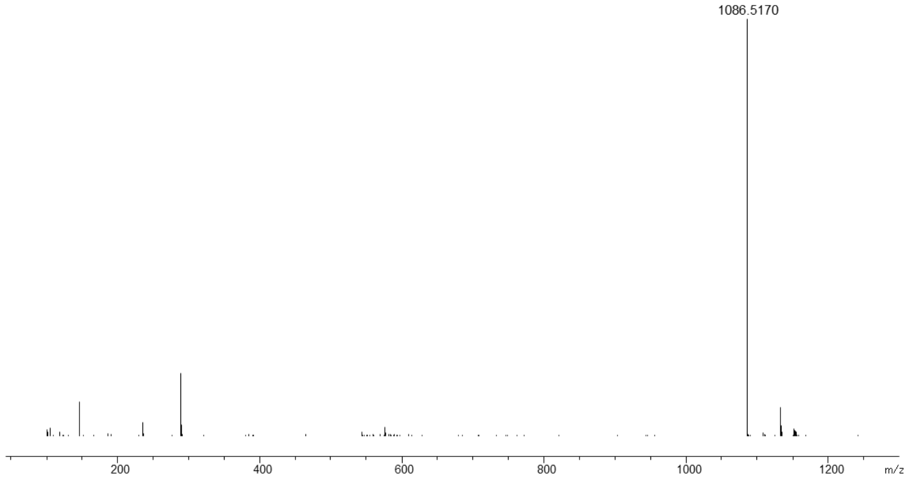

#### Synthesis of MP3

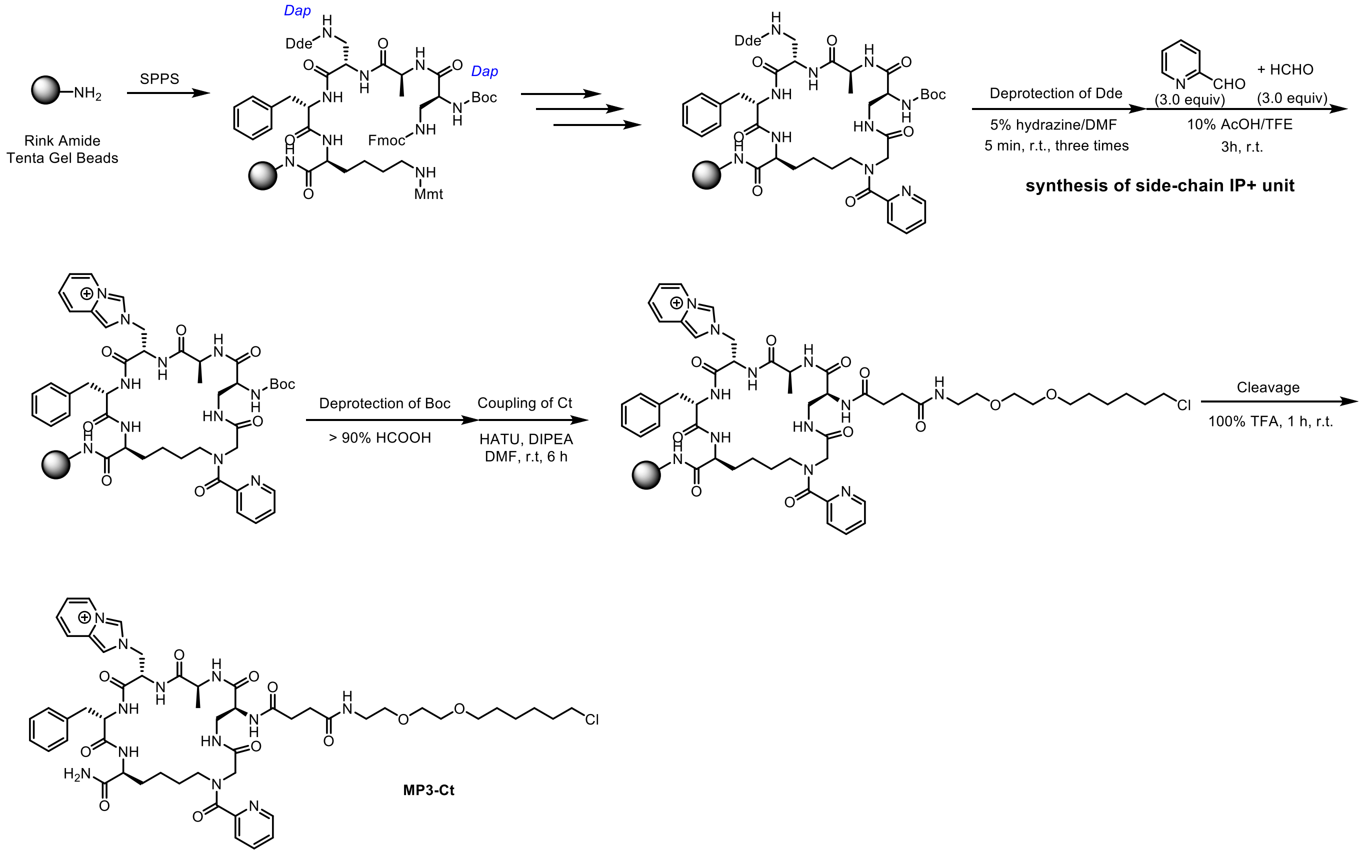


**Supplementary Scheme 3.** Synthetic route to **MP3-Ct**. Preparation of macrocycle with side-chain IP$^{+}$ unit

Purified HPLC (210 nm) and HRMS of MP3-Ct:

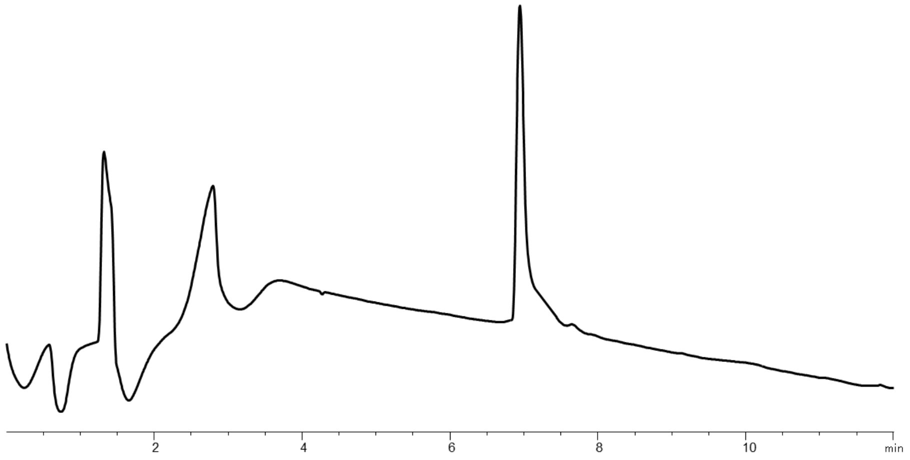


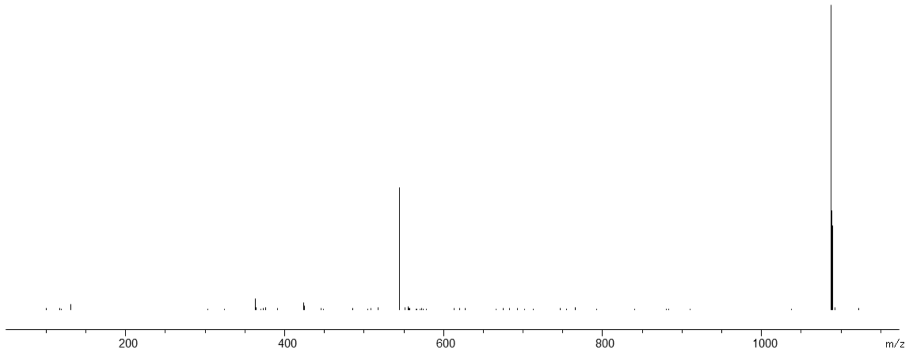

#### Synthesis of MP4

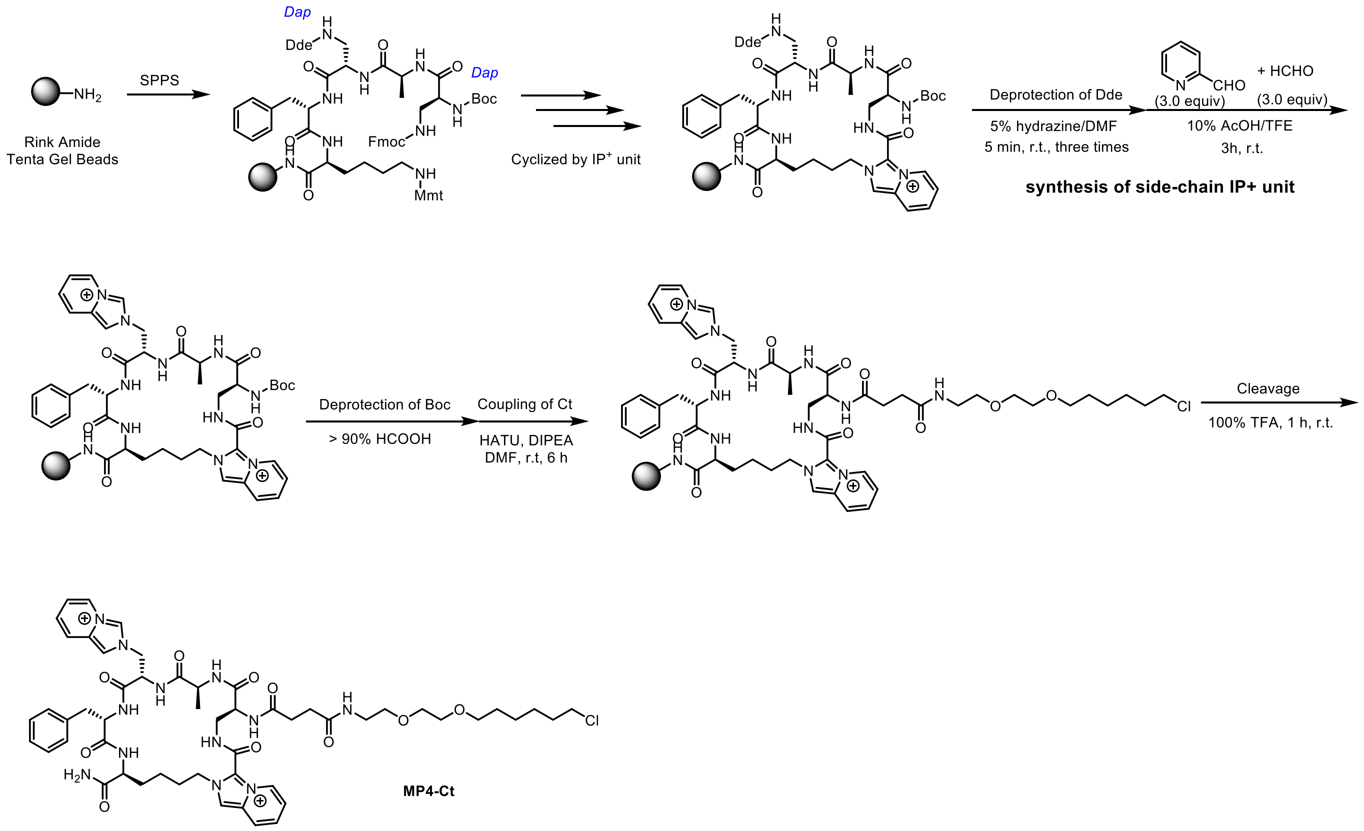


**Supplementary Scheme 4.** Synthetic route to MP4-Ct. Preparation of macrocyclic peptide containing two $IP^+$ units

Purified HPLC (210 nm) and HRMS of MP4-Ct:

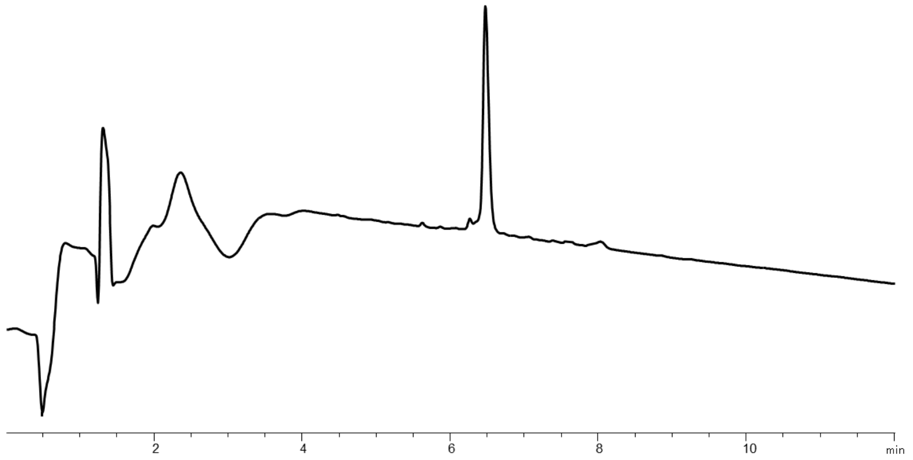


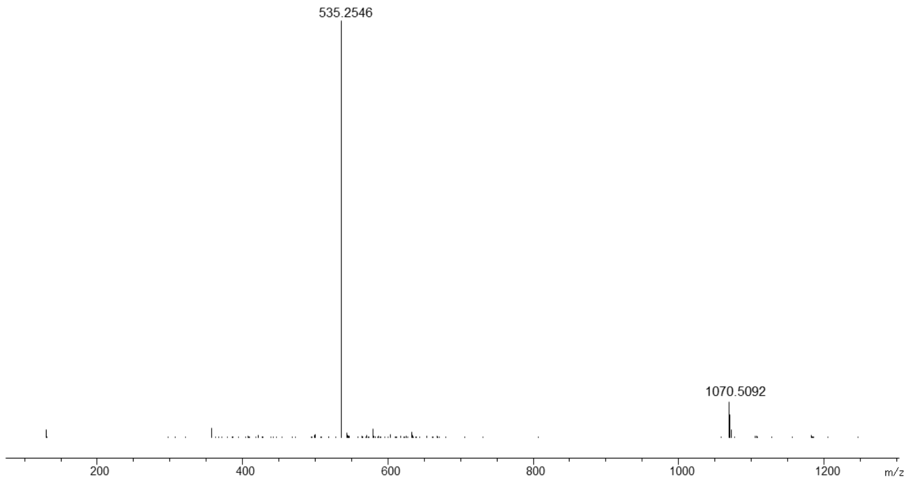

#### Synthesis of MP5

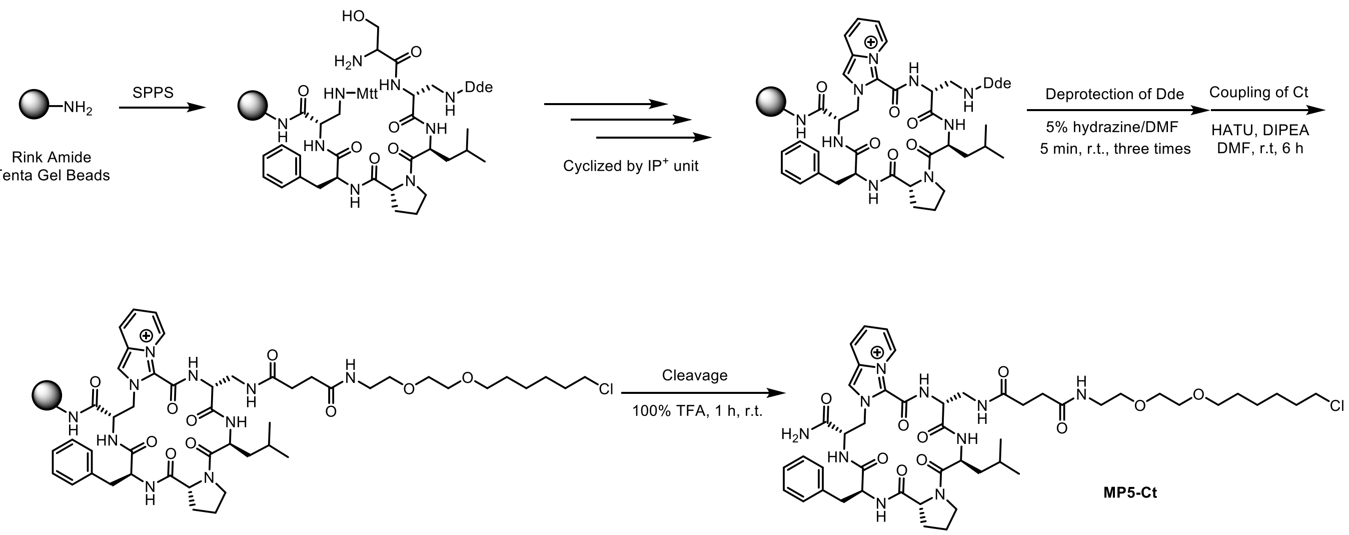


**Supplementary Scheme 5.** Synthetic route to MP5-Ct.

Purified HPLC (210 nm) and HRMS of MP5-Ct:

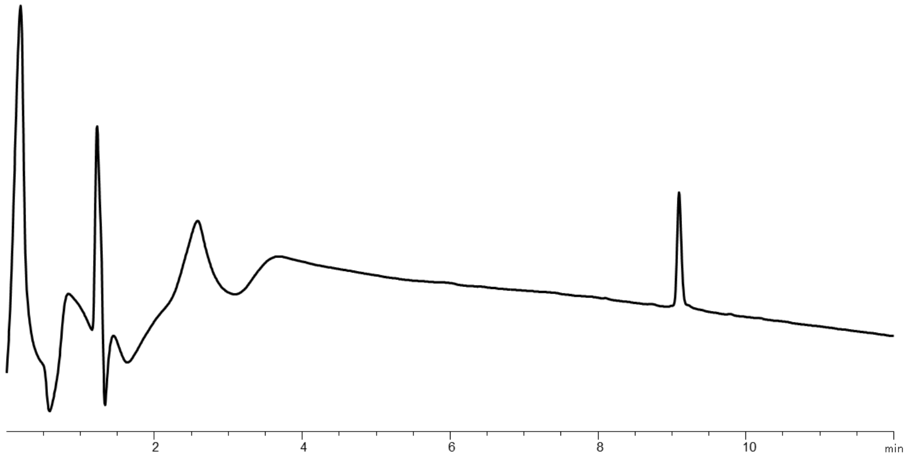


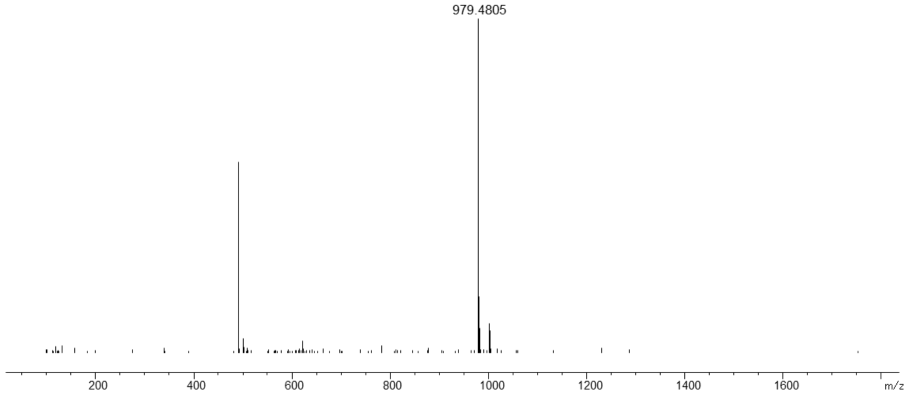

#### Synthesis of MP6

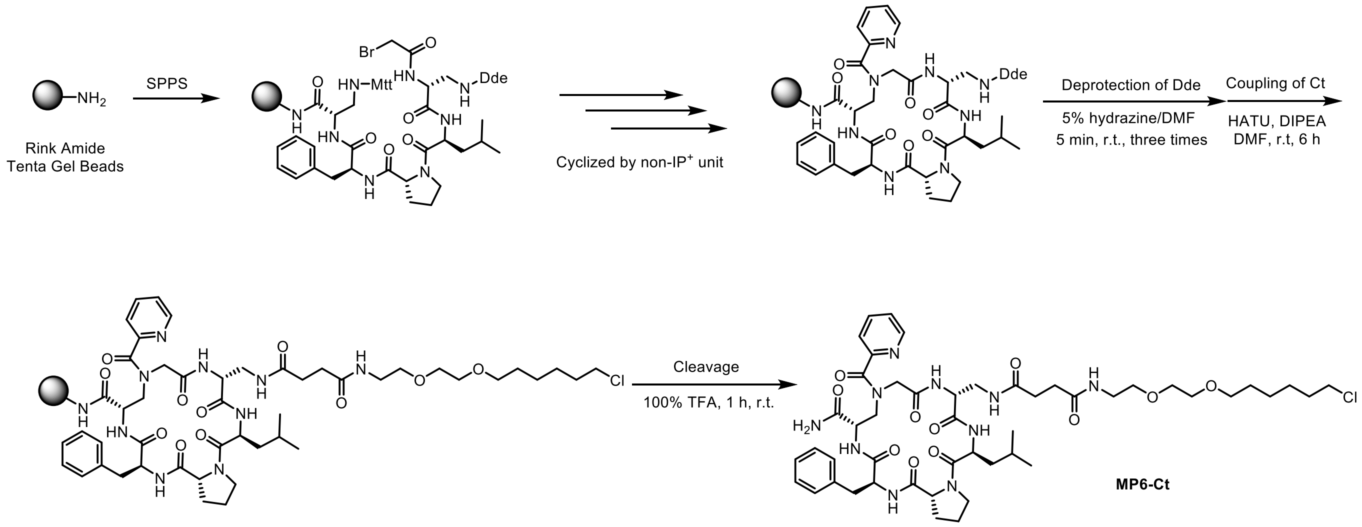


**Supplementary Scheme 6.** Synthetic route to MP6-Ct.

Purified HPLC (210 nm) and HRMS of MP6-Ct:

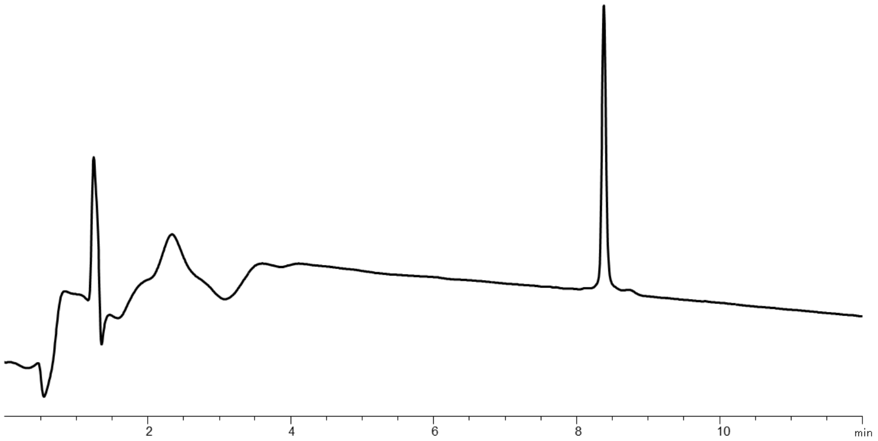


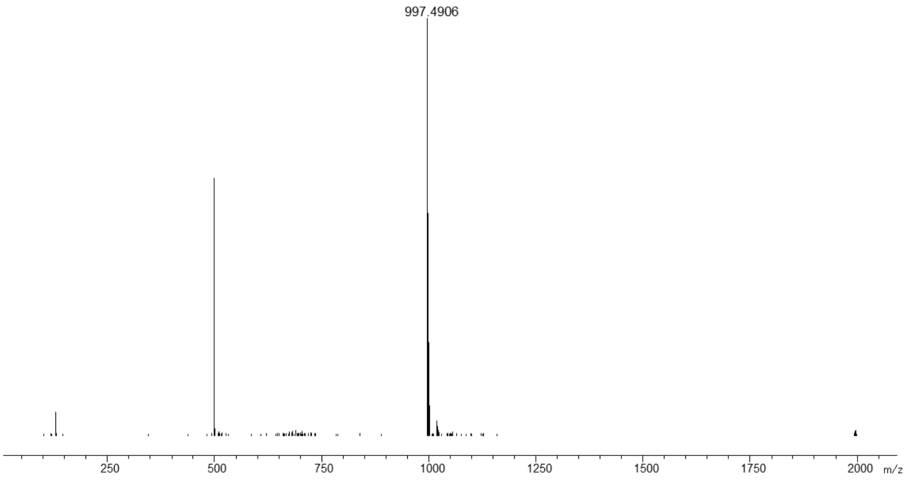

### Synthesis of MP7

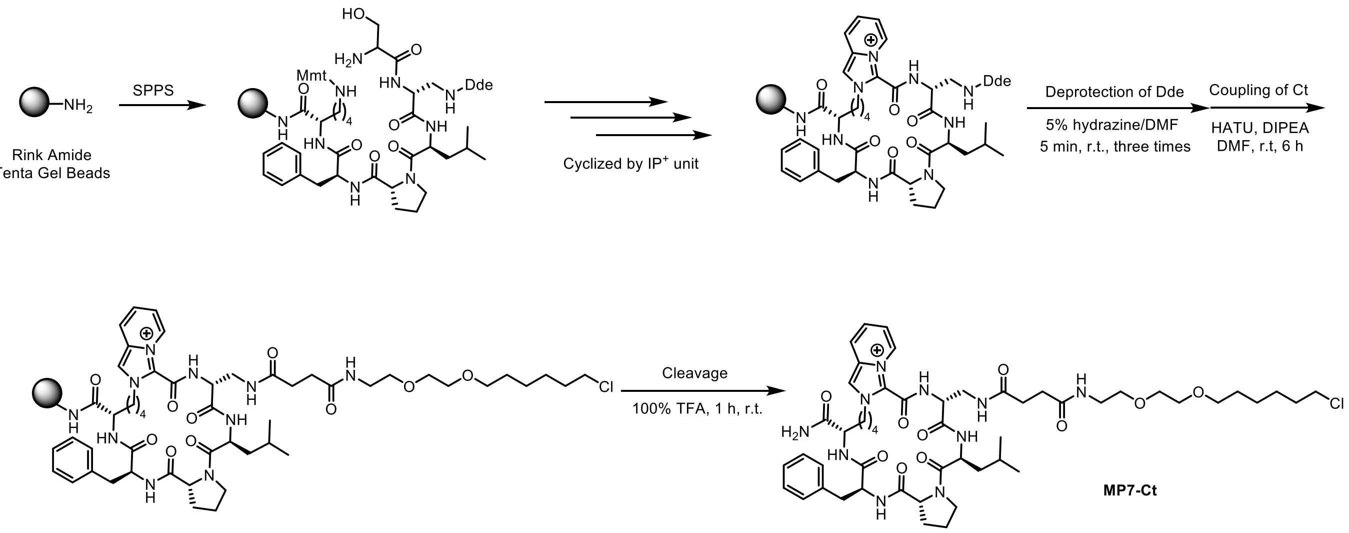


**Supplementary Scheme 7.** Synthetic route to MP7-Ct.

Purified HPLC (210 nm) and HRMS of MP7-Ct:

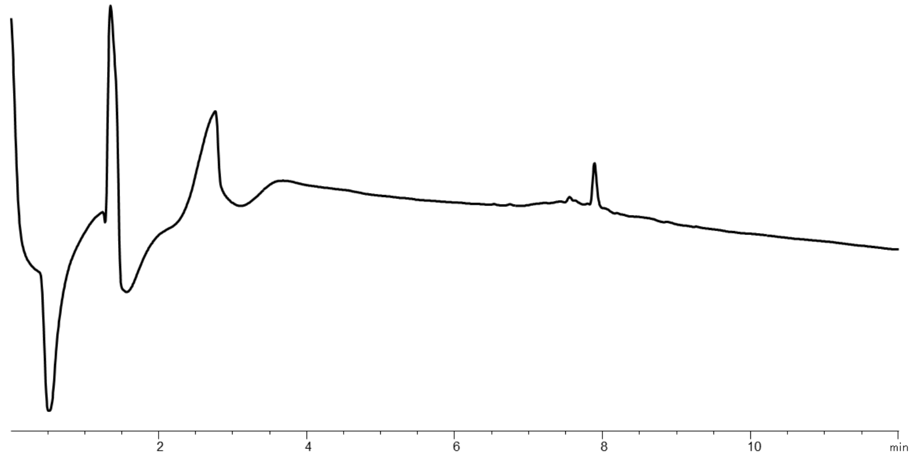


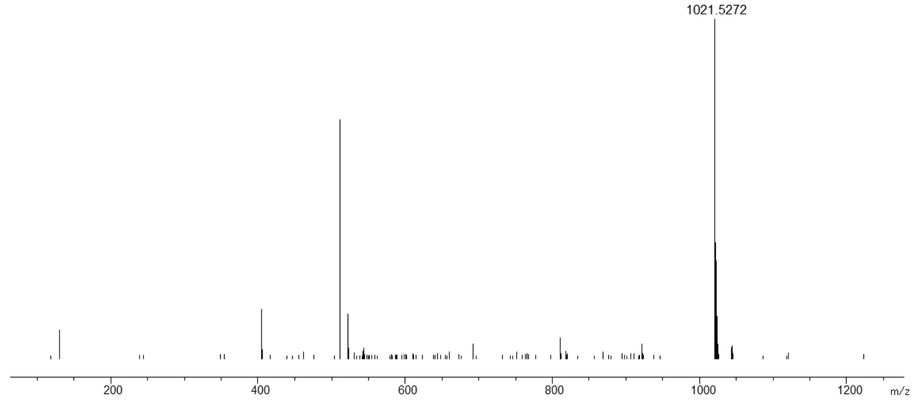

### Synthesis of MP8

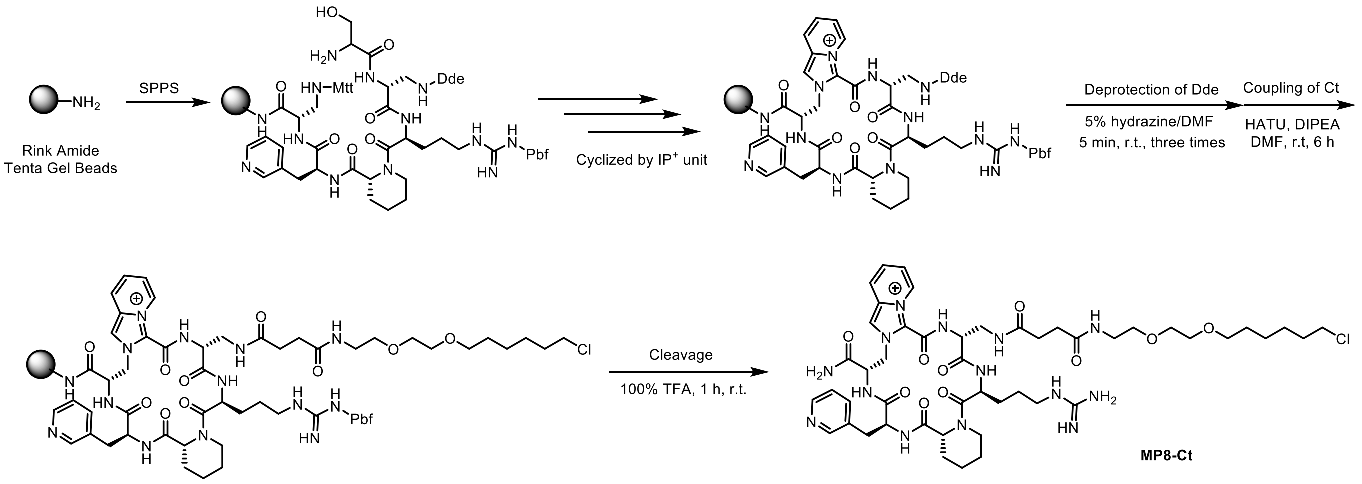


**Supplementary Scheme 8.** Synthetic route to MP8-Ct.

Purified HPLC (210 nm) and HRMS of MP8-Ct:

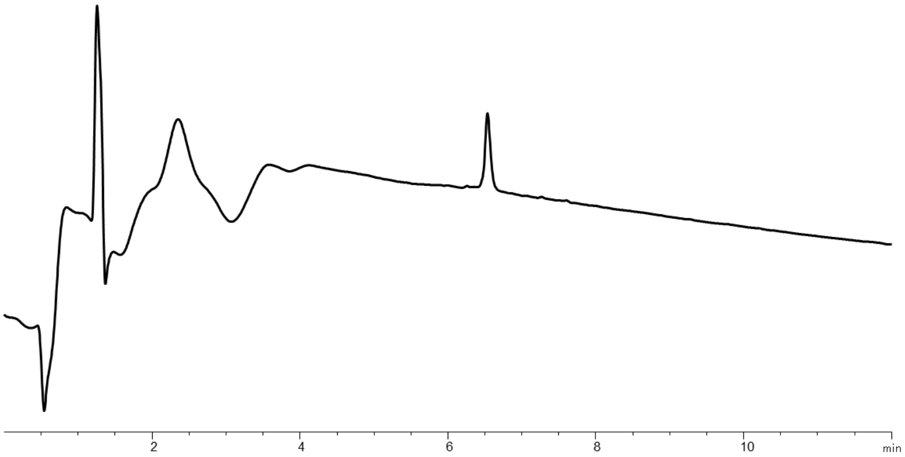


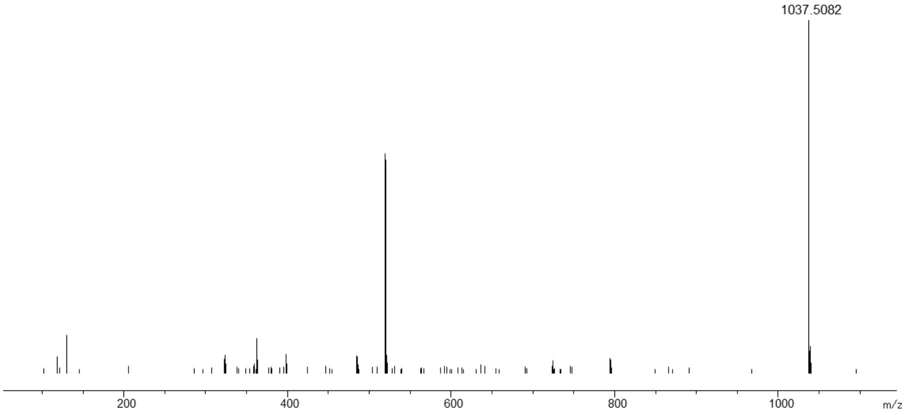

### Synthesis of MP9

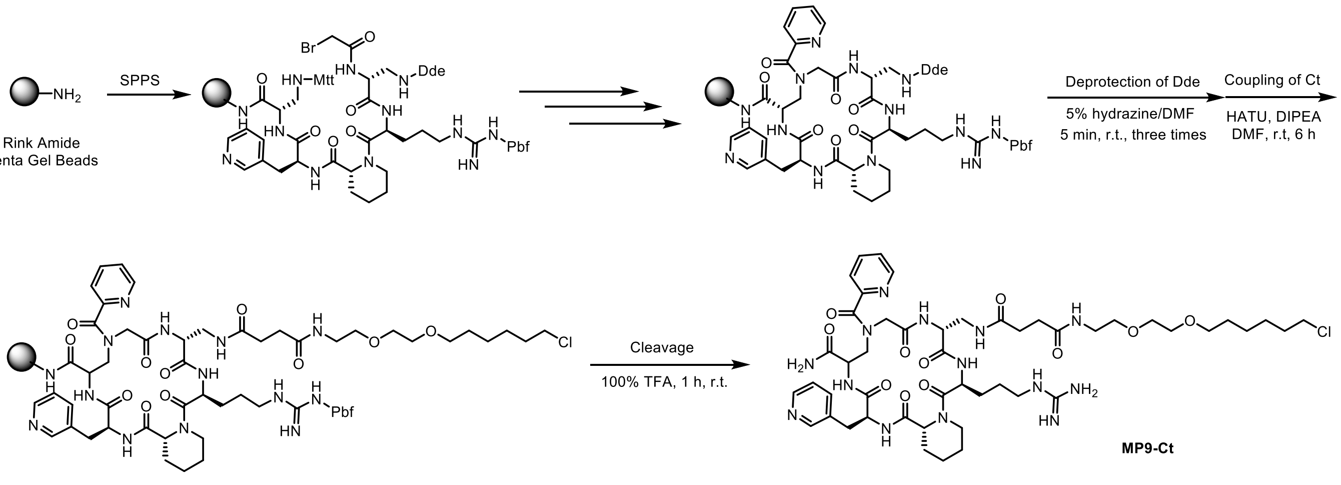


**Supplementary Scheme 9.** Synthetic route to MP9-Ct.

Purified HPLC (210 nm) and HRMS of MP9-Ct:

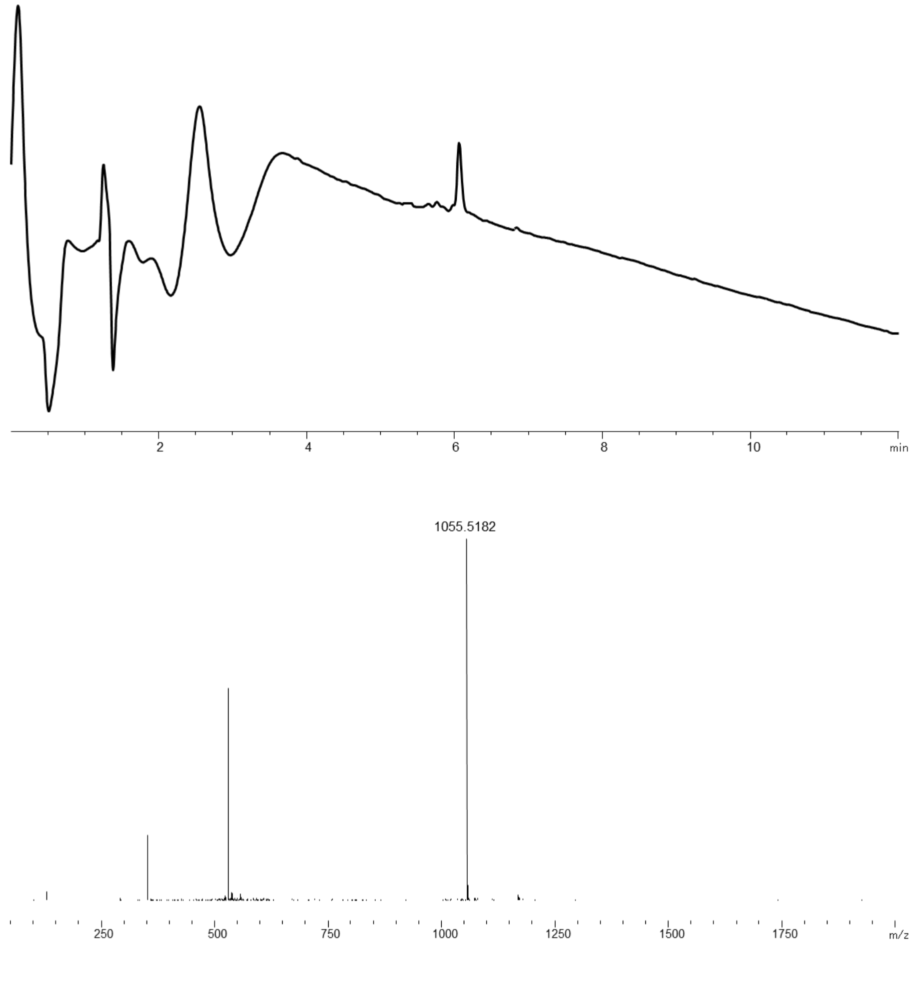

### Synthesis of MP10

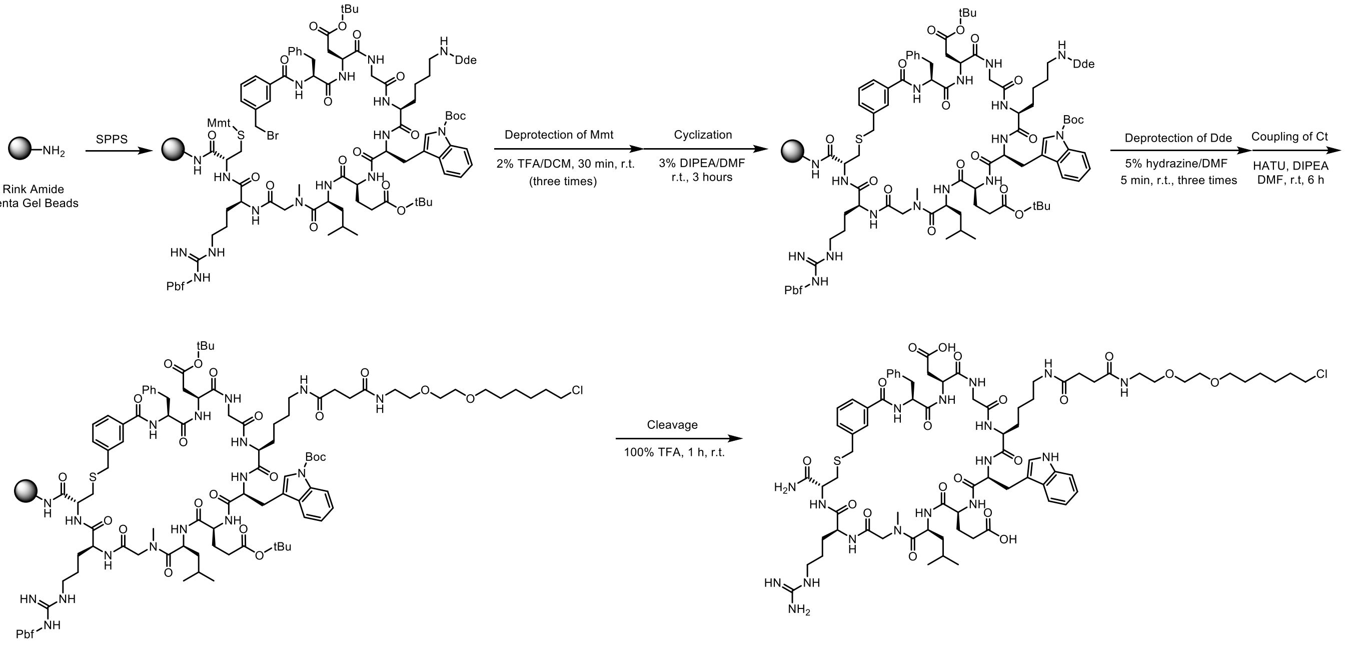


**Supplementary Scheme 10.** Synthetic route to MP10-Ct.

Purified HPLC (210 nm) and HRMS of MP10-Ct:

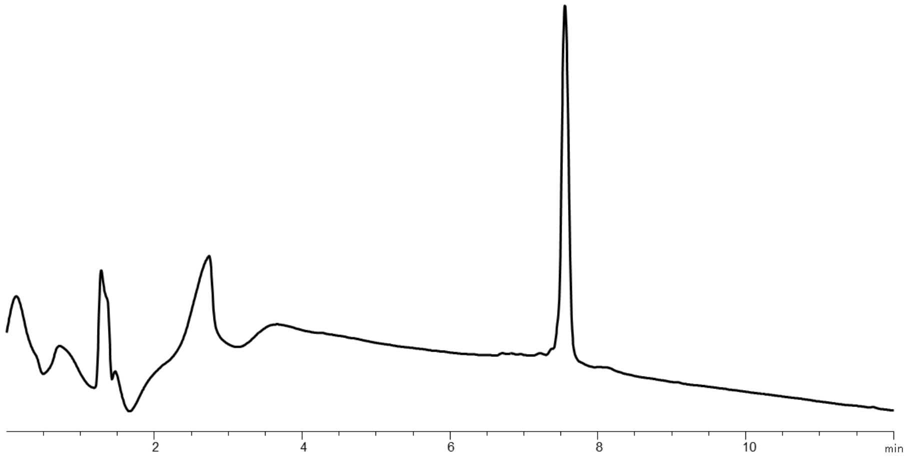


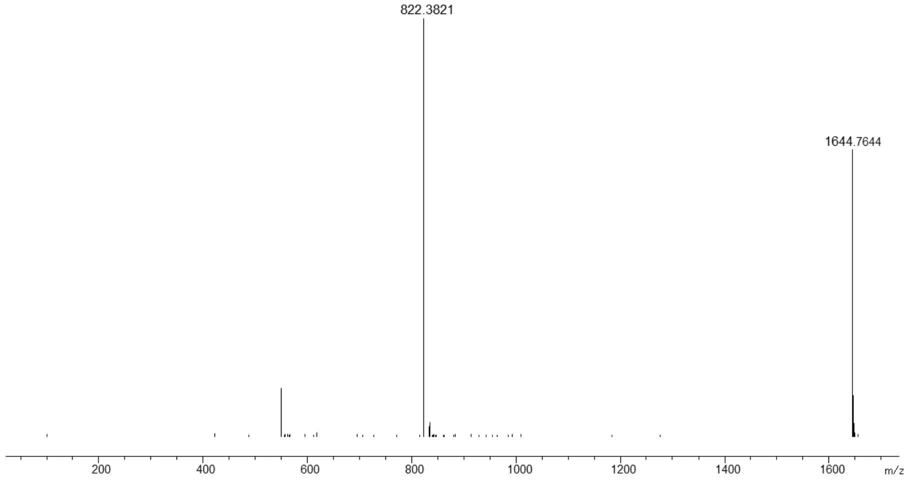

### Synthesis of MP11

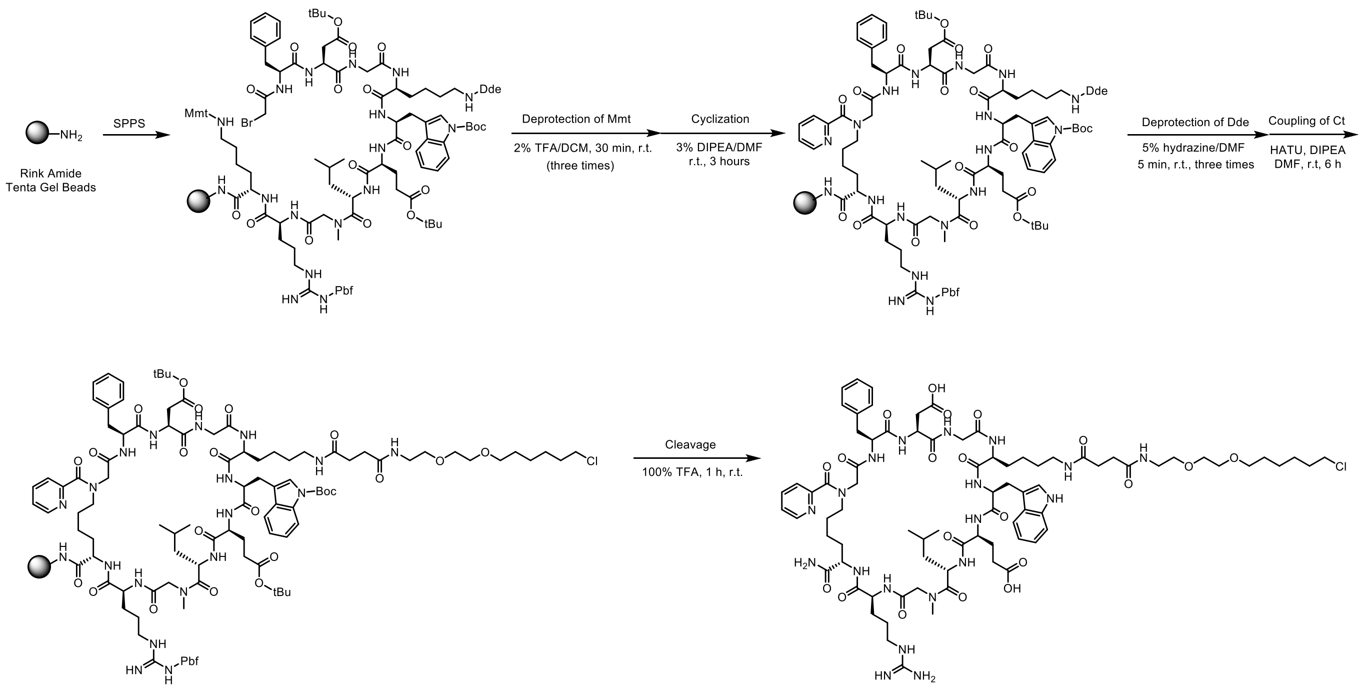


**Supplementary Scheme 11.** Synthetic route to MP11-Ct.

Purified HPLC (210 nm) and HRMS of MP11-Ct:

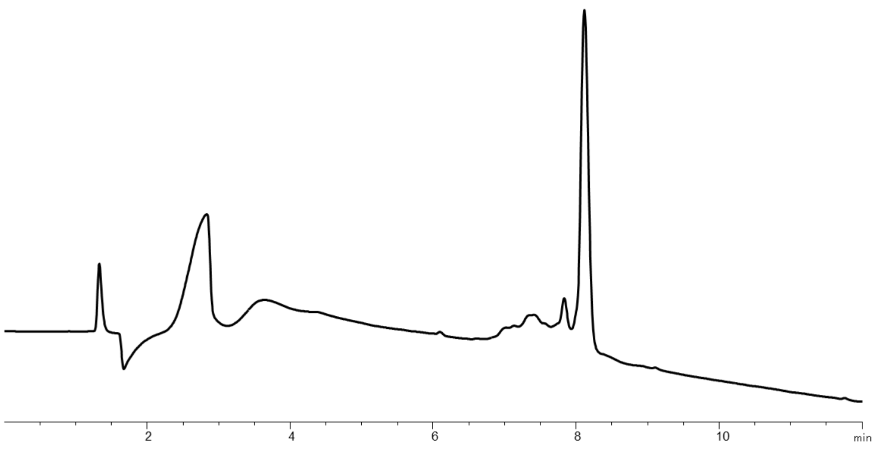


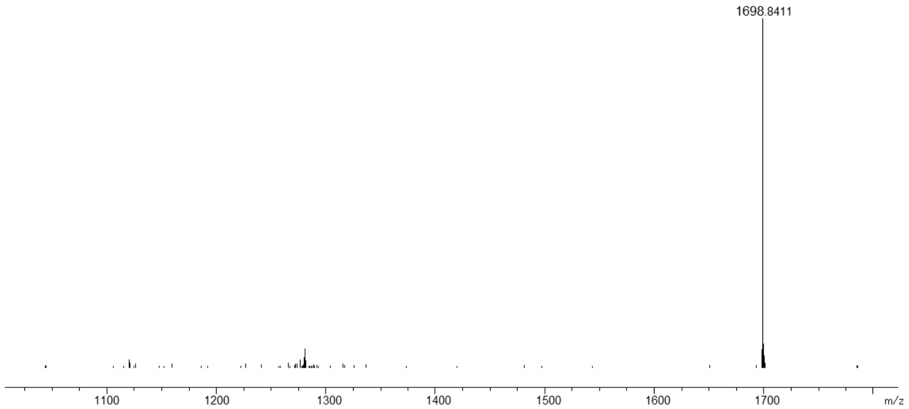

### Synthesis of MP12

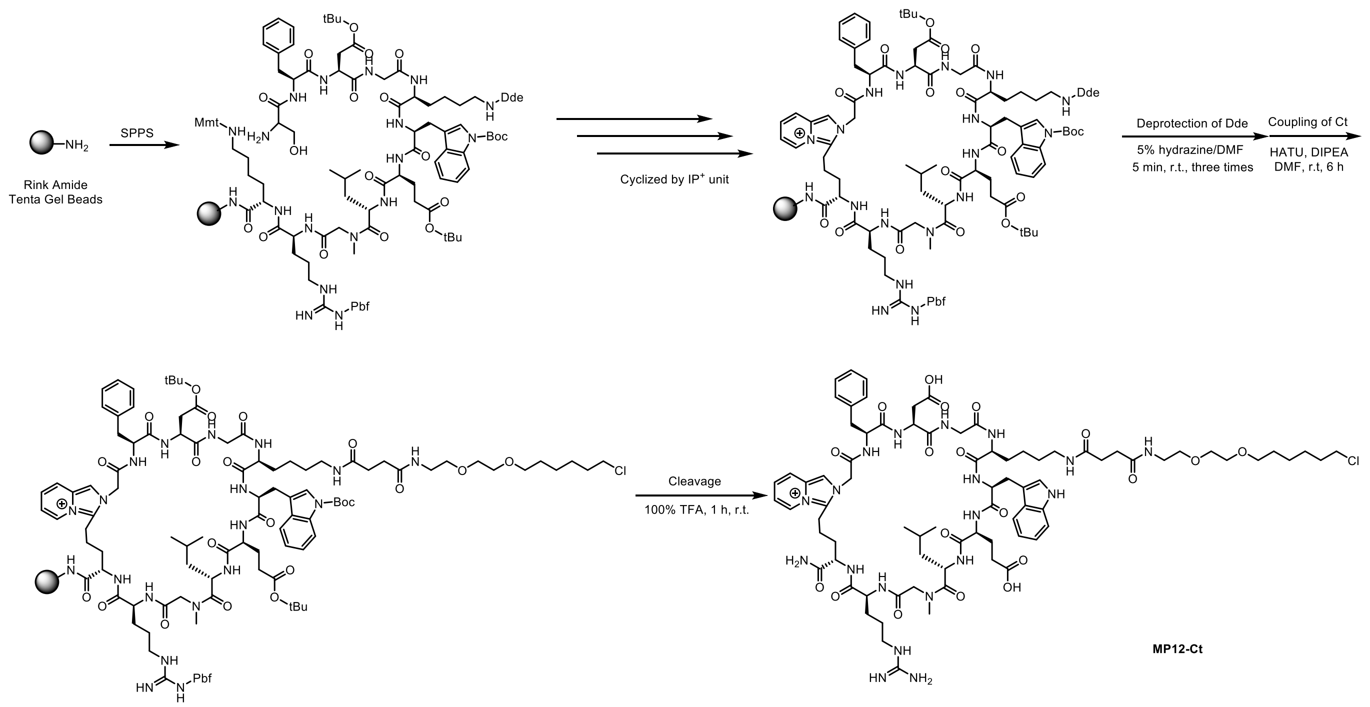


**Supplementary Scheme 12.** Synthetic route to MP12-Ct.

### Synthesis of MP13

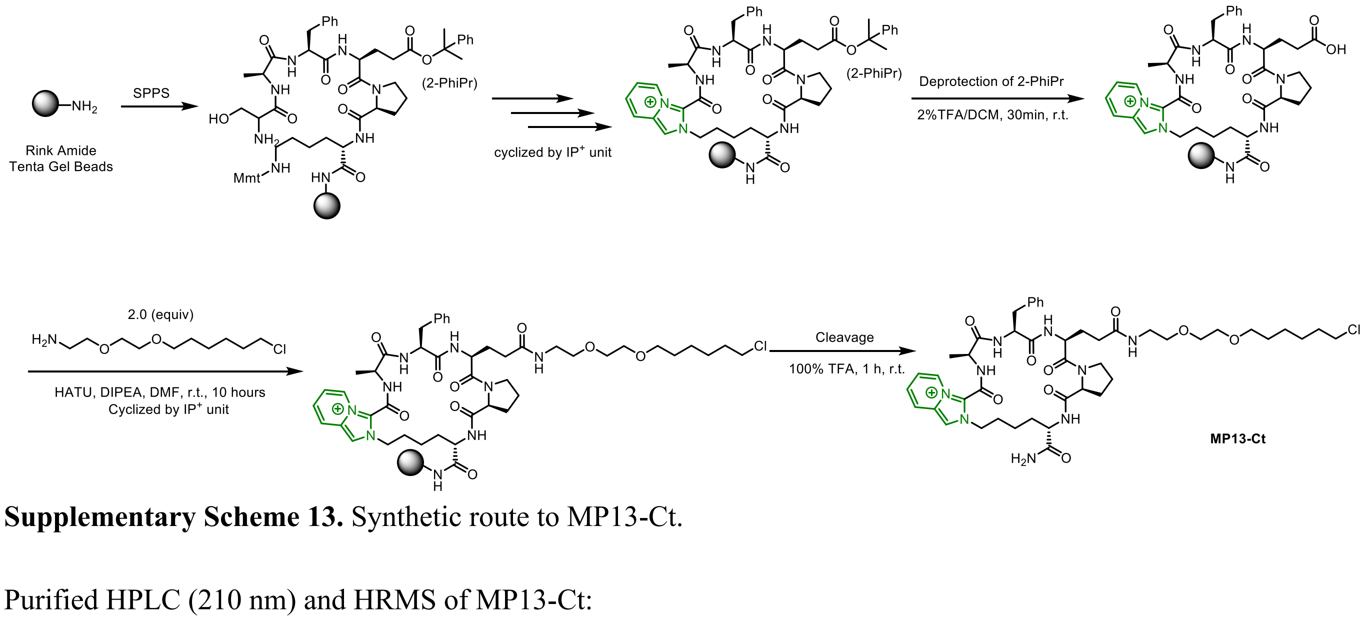


**Supplementary Scheme 13.** Synthetic route to MP13-Ct.

Purified HPLC (210 nm) and HRMS of MP13-Ct:

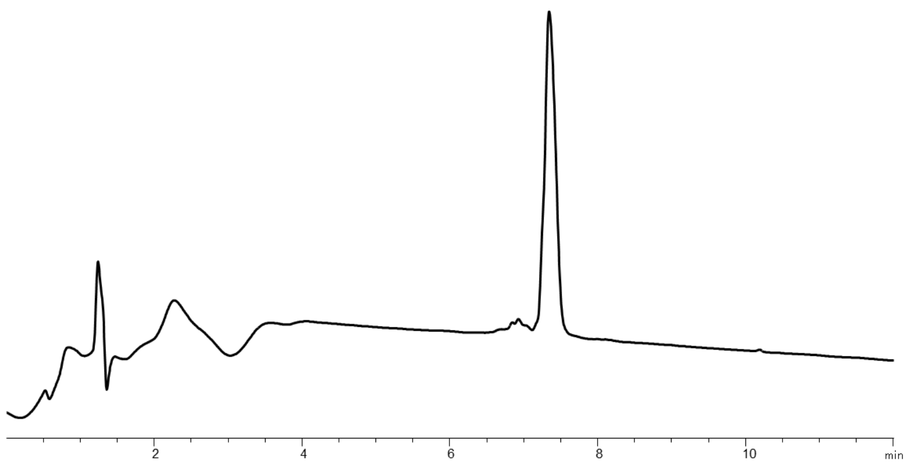


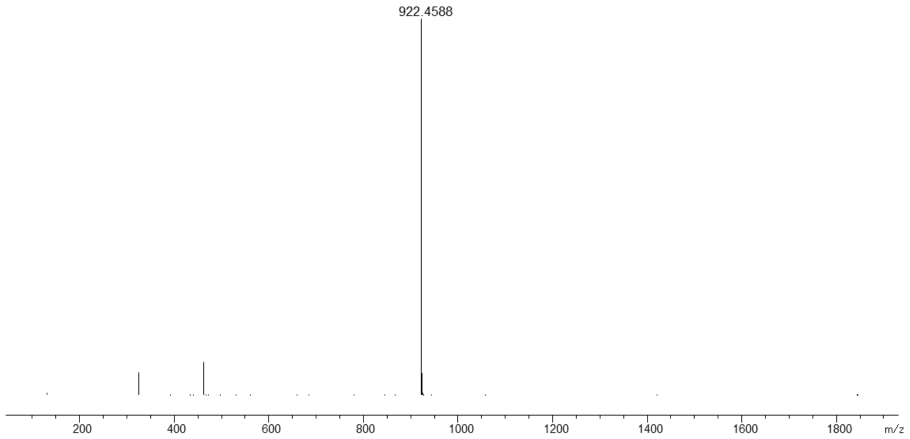


### Synthesis of MP14

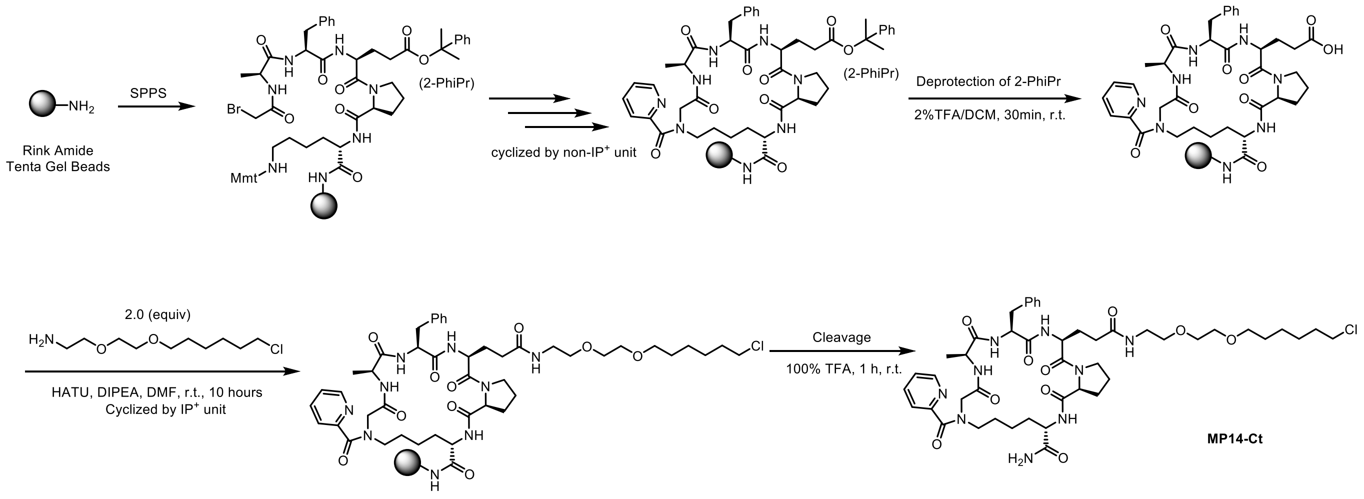


**Supplementary Scheme 14.** Synthetic route to MP14-Ct.

Purified HPLC (210 nm) and HRMS of MP14-Ct:

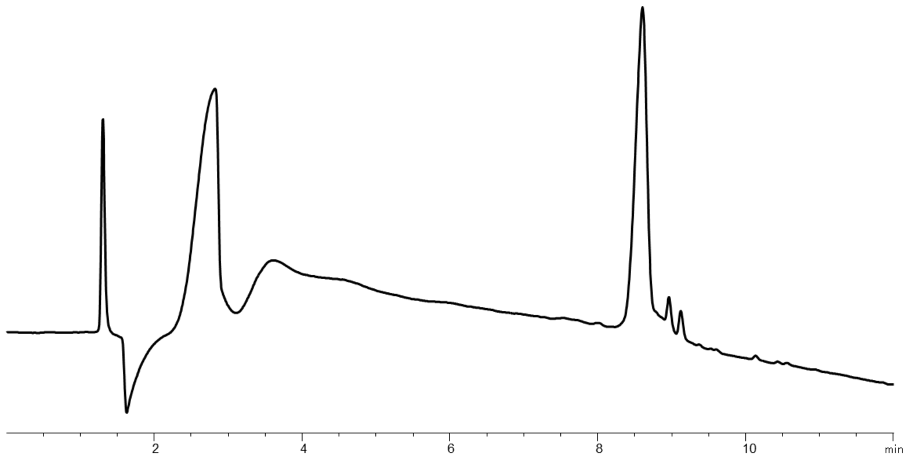


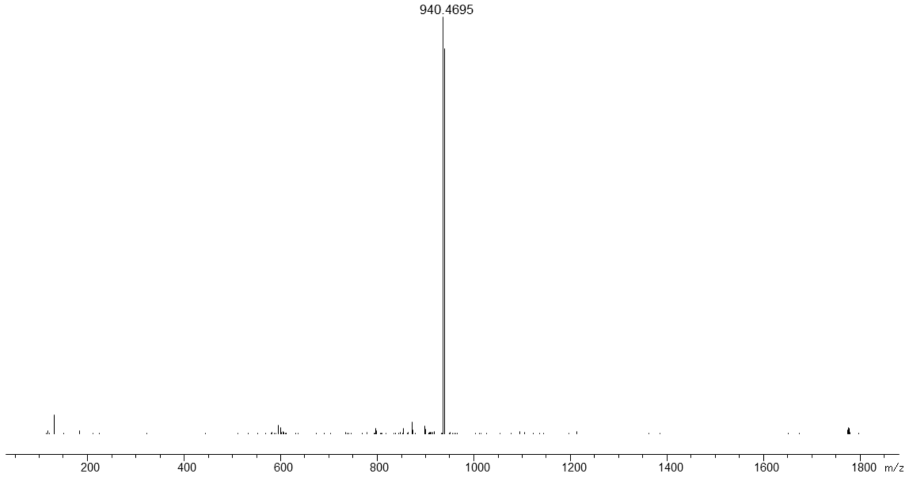


#### Synthesis of UNP-6457, MP23 and MP23-L

**UNP-6457** was prepared according to the reported procedure (ACS Med. Chem. Lett. 2023, 14, 820).

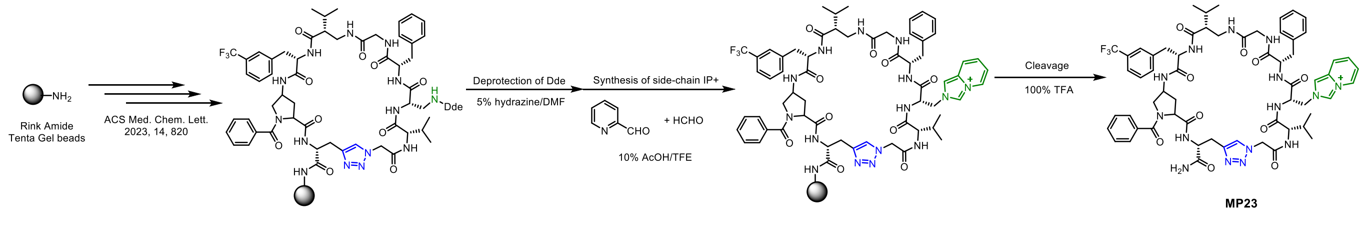


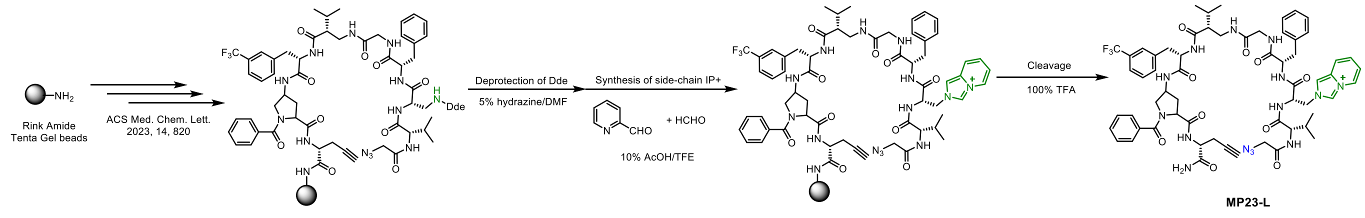

Purified HPLC (210 nm) and HRMS of **UNP-6457**:

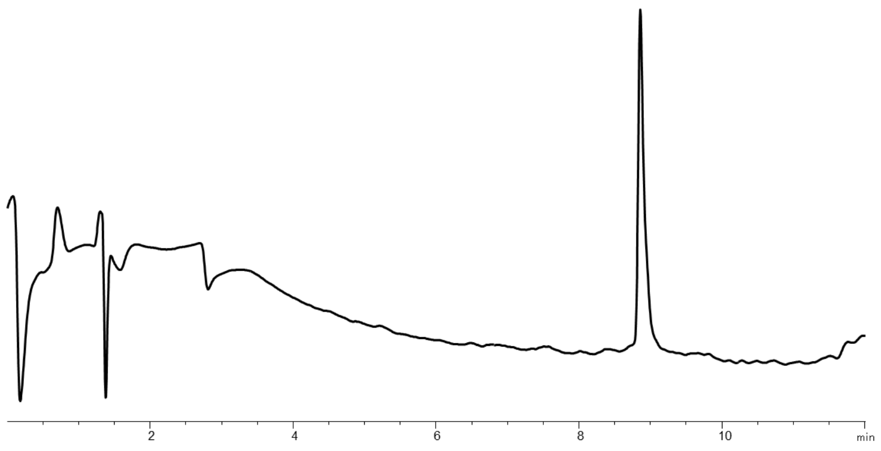


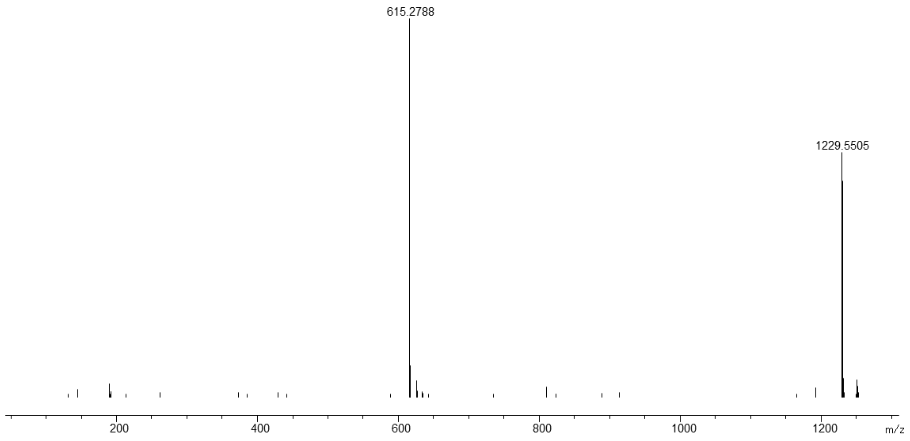


Purified HPLC (210 nm) and HRMS of **MP23**:

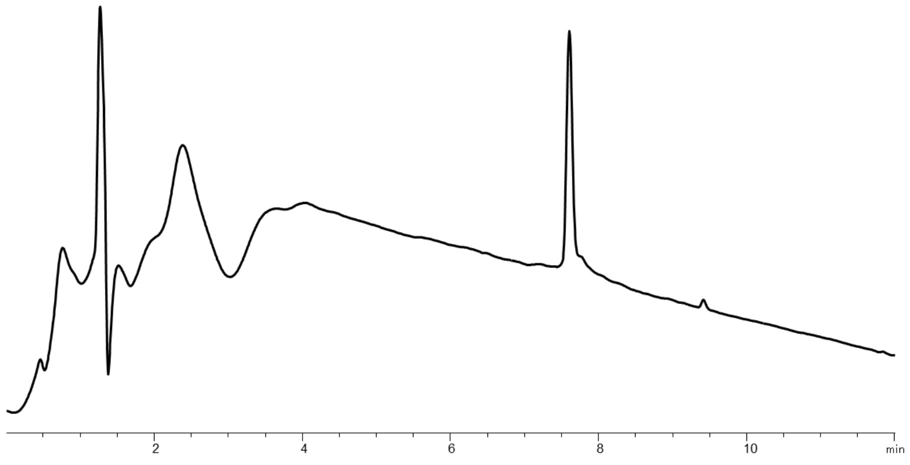

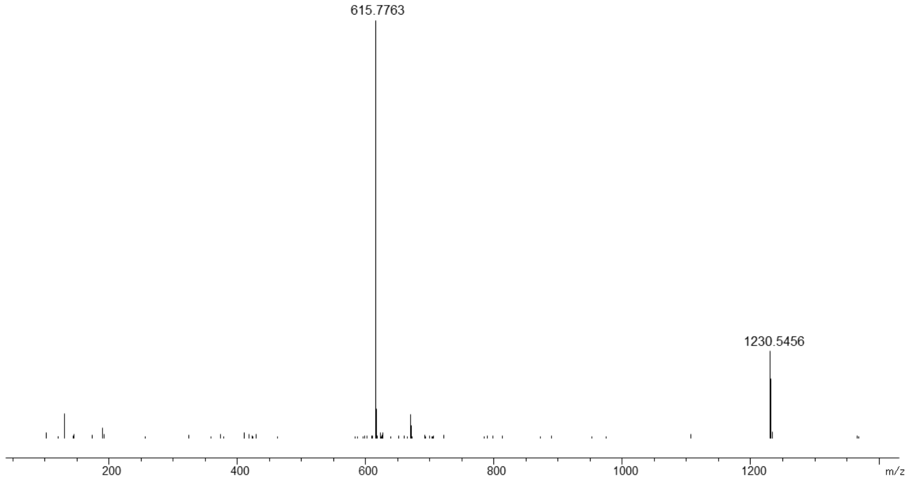


Purified HPLC (210 nm) and HRMS of **MP23-L**:

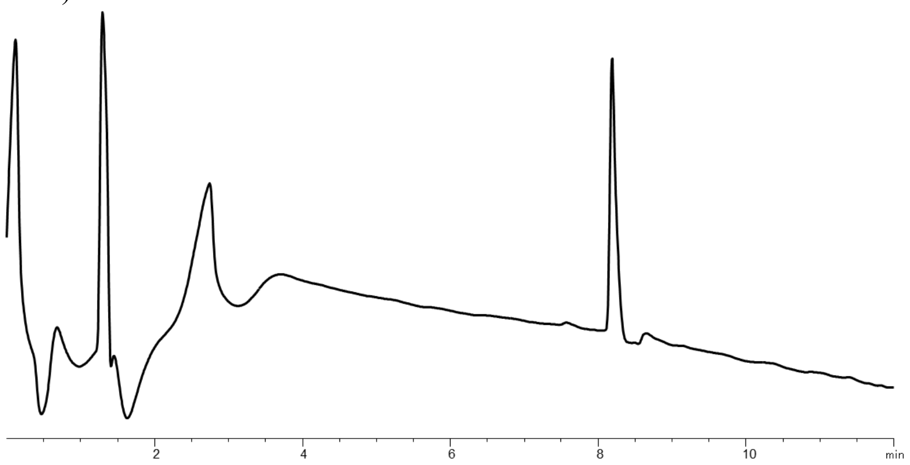


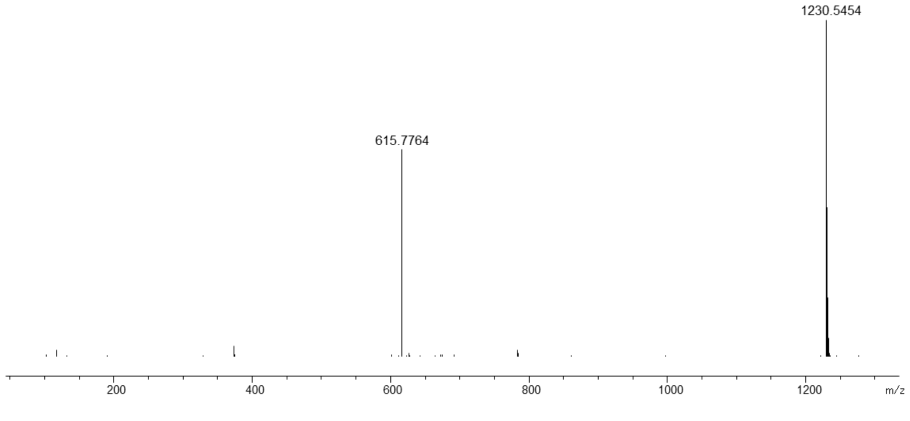

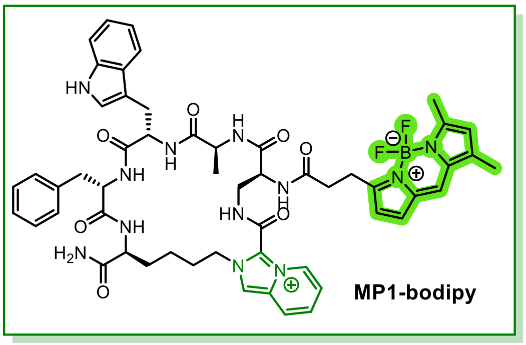


Purified HPLC (210 nm) and HRMS of **MP1-bodipy**:

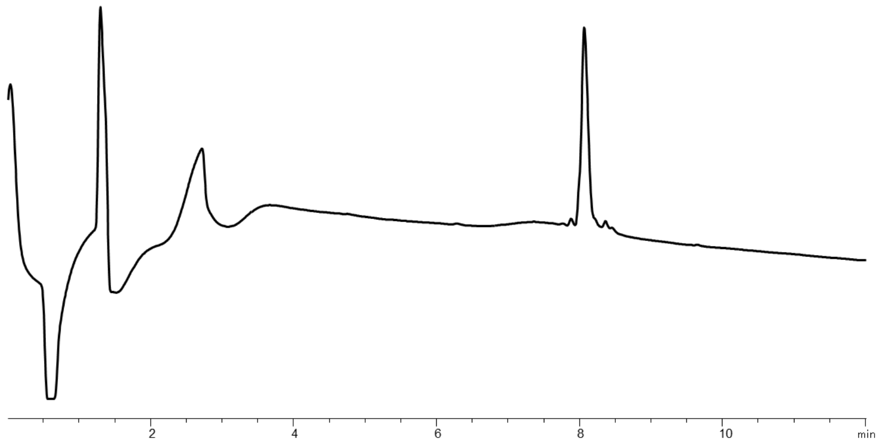


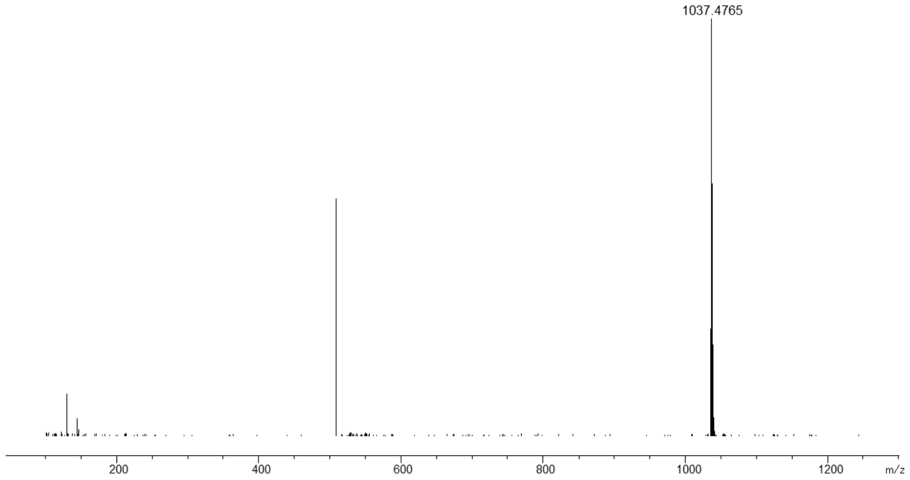

#### Chloroalkane Penetration Assay (CAPA) of IP$^{+}$-linked MPs

HEK293 cells expressing HaloTag7 were seeded in 24 well plates. The next day, cells were treated with ct-MPs and incubated at 37°C for four hours. After incubation, cells were washed then incubated with 5µM Tamra-ct in DPBS (Gibco 14190-144) supplemented with 1% bovine serum albumin BSA for 30 min at 4°C. Cells were again washed, resuspended in 1% BSA in DPBS, and fluorescence was analyzed on a BD LSR II Flow Cytometer. Data were subsequently analyzed on FlowJo and normalized to a DMSO-only positive control. Tamra-ct was prepared according to the previously reported procedure. (J. Kritzer. *et al.* J. Am. Chem. Soc. **2018**, *140.* 11360).

#### Membrane pore experimental protocol

HEK293 cells were seeded overnight in a 24 well plate. In the morning, cells were incubated for 2 hours at 37°C with 1uM propidium iodide and either 10µM MP1 or 10µM MP2. Cells pre-treated for 30 minutes with 0.2% Tween-20 served as a positive control and DMSO-treated cells served as a negative control. Following incubation, cells were washed, resuspended in PBST and propidium iodide staining was determined on a BD LSR II Flow Cytometer. Data were subsequently analyzed on FlowJo.

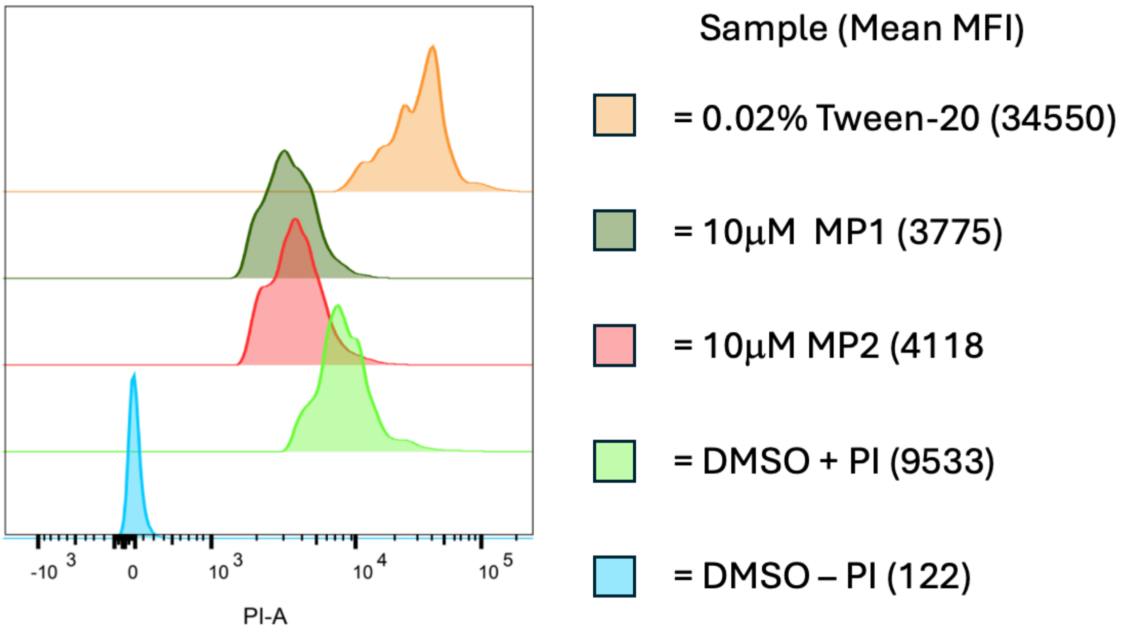


**Supplementary Scheme 15.** Membrane pore experimental data

#### Imaging study method

Cells were plated onto glass coverslips in 12-well plates, then treated with MPs at indicated concentrations. After the indicated time of exposure, the media was removed, cells were washed 1x in DPBS and then fixed in 500uL 4% paraformaldehyde for 12 minutes. To permeabilize the cells, pfa was then removed, and 1mL 75% ethanol was added. Cells were then stored at 4ºC overnight before staining. For staining of immunofluorescence, cells were washed 2x in PBS, then incubated in 500µL primary antibody in PBS for four hours at 4ºC. Cells were washed 2x in PBS, then secondary antibodies were added for two hours at 4ºC. Cells were washed 2x, then fixed in 4% pfa for 10 minutes. Slips were then mounted on slides (Fisherbrand 12-544-1) with Vectashield (VectorLabs H-1200) and dried.

All Images were taken on a Nikon Eclipse Ti2 confocal microscope with a 100x, 1.45 NA oil objective or a 20x, NA objective. The filter set included: C-FL DAPI Filter Set, High-Signal-Noise, Semrock Brightline®, Excitation: 356/30nm (341-371nm). Super-resolution microscopy was imaged using GATACA Systems Live-SR. For immunofluorescence detection, cells were incubated with Rabbit mAb anti-Rab7 antibody (D95F2) (Cell Signaling Technology: 9367) (1:500), Mouse mAb anti-E-Cadherin (4A2) (Cell Signaling Technology: 14472) (1:500), and Rabbit polyclonal MAVS Antibody (Cell Signaling Technology: 3993) (1:200) primary antibodies at 4C overnight, washed three times, and then incubated with Goat Anti-Rabbit IgG H&L (Alexa Fluor® 647) (Abcam: ab150079) and Goat Anti-Mouse IgG H&L (FITC) (Abcam; ab97022) at 1:1000 for two hours. Live cell imaging was completed using Cellvis glass bottom dishes (D35C4-20-0-N). Cells were seeded at 80% confluency and imaged while kept at 37°C at 5% $CO_2$ in microscope incubation box. Cells were treated as indicated and imaged at indicated intervals. Mitochondria stained with MitoTracker® Deep Red (CST #8778).

Image Analysis

Images were processed using Nikon NIS-Elements Advanced Research software and Fiji ImageJ image processing software package. Images use indicated single slice or stacked Z planes (stacked) and max/min display values were adjusted to properly view fluorescence intensity for each channel. Macros and built-in functions were used to analyze images for indicated variables. ImageJ macros and CellProfiler pipelines available upon request.
